# SOUR Disentangles Heritable and Microenvironmental Contributions to Intratumoral Heterogeneity in Spatial Lineage Tracing Data

**DOI:** 10.64898/2026.11.19.767955

**Authors:** Eric Weine, David Burt, Karan Joshua Abraham, William Colgan, Jonathan Weissman, Rafael A. Irizarry, Tamara Broderick, Joshua Schraiber

## Abstract

Intratumoral heterogeneity is hypothesized to be a major contributor to treatment resistance, yet the processes that generate it remain only partially understood. New technologies enable measurement of gene expression in individual tumor cells while also determining where the cells are located within a tumor and how they are related through cell division. A central question for such data is how much gene expression differences reflect characteristics passed from parent cells to their descendants, rather than influences from their local surroundings. Distinguishing these influences is challenging because closely related cells often reside near one another. Existing methods examine the relationship between gene expression and each of these factors separately, and can therefore incorrectly attribute spatial effects to ancestry or inherited effects to spatial location. Here, we present SOUR, a statistical method that uses both cell ancestry and spatial location to estimate their respective contributions to differences in gene expression. Using simulated data and two spatial lineage tracing datasets, we demonstrate that SOUR can estimate the separate contributions of cell ancestry and spatial location to variation in gene expression, while substantially reducing the confounding observed with existing approaches.

## 1 Background

Intratumoral heterogeneity (ITH), or the presence of diverse molecular states among cells within a single tumor, is a common feature of human cancers [1, 2]. ITH is hypothesized to be a major contributor to treatment resistance, as no single treatment can target the spectrum of cellular states present in a tumor. Understanding what drives this heterogeneity and how it evolves as a tumor grows has consequently become a major goal of cancer research [3, 4].

Broadly, ITH is thought to arise from two sources. First, as tumor cells divide, they accumulate genetic and epigenetic alterations, including changes in copy number [5] and DNA methylation [6], which alter gene expression and are passed on to progeny. These alterations may arise spontaneously within a cell, but they may also be induced by intercellular signaling. We refer to such effects as *heritable variation*. Second, cells adopt transient states in response to the niche they currently occupy. The tumor microenvironment is complex and dynamic, and cells in different locations experience different conditions, including the availability of oxygen and nutrients and the composition of neighboring stromal and immune cells. The tumor core, for example, often has limited access to nutrients and becomes hypoxic [7]. We refer to such effects as *microenvironmental regulation*.

Recently developed technologies have allowed biologists to identify both spatial and ancestral patterns of gene expression in model systems and patient tumors. Spatial transcriptomics technologies have been used to map ITH across tissue regions and to examine how microenvironmental interactions may shape tumor cell states [8–12]. In parallel, lineage tracing technologies have been used to reconstruct tumor phylogenies and to characterize how gene expression changes as tumors evolve [13–17]. However, spatial and lineage distances between cells are often highly correlated, so measuring only one of these variables in an experiment limits insight into the mechanisms driving ITH. Heritable variation may appear to result from microenvironmental regulation because tumor clones are typically confined to limited spatial regions [18, 19]. Conversely, microenvironmental regulation may appear heritable when particular environmental conditions affect only a small, spatially confined set of clones.

Motivated in part by these limitations, recently developed technologies can *simultaneously* profile lineage, spatial, and transcriptomic information in a single experiment [20–27]. These data raise a natural question: to what extent is transcriptional heterogeneity within a tumor associated with microenvironmental forces rather than heritable variation? Indeed, a central motivation for developing these technologies is that previous approaches “limited our ability to parse the relative contributions of nature versus nurture in governing cell behavior” [28].

However, currently used data analysis approaches for assessing the contributions of microenvironmental and heritable forces do not take full advantage of the spatial and ancestry information measured for each cell. For instance, the analysis in [23] calculates the spread of mean expression across clones and, separately, across spatial regions to obtain a *clone score* and a *space score*. Simi-larly, the analysis in [21] calculates gene expression autocorrelation with respect to lineage distance and spatial distance separately. Both approaches are therefore subject to confounding: expression variation associated solely with microenvironmental forces can score highly on measures of heritable variation, and vice versa.

Similar challenges have been encountered in comparative phylogenetics, where biologists are interested in understanding the effects of ancestry and environmental factors on the evolution of quantitative phenotypes among related species. Because closely related species tend to occupy similar environments, analyzing each variable in isolation can lead to an over-estimate of its importance. Freckleton & Jetz addressed this problem by developing a random effects regression model that *simultaneously* estimates the effects of ancestry and habitat on a phenotype [29]. Here, we build on their approach to disentangle heritable and microenvironmental contributions to gene expression variation in single-cell spatial lineage tracing data. Applying this approach to cells requires adapting the model to account for two key features of these data. First, Freckleton & Jetz assume that traits are normally distributed, whereas gene expression counts measured by spatial lineage tracing technologies are sparse and heteroskedastic, and are better modeled by count-based distributions [30, 31]. Second, and more fundamentally, Freckleton & Jetz assume that spatial covariance between the traits of two species arises solely from the environments their common ancestors experienced. Under this model, two distantly related species will show little spatial covariance between their traits even if they occupy the same environment today. While this assumption may be reasonable in the context of modeling species traits that evolve over millions of years, it is poorly suited for accounting for the effects of cellular microenvironments that can rapidly induce similar phenotypes in completely unrelated cells through cell signaling [32, 33].

The presence of distantly related cells in shared microenvironments also provides an opportunity to distinguish microenvironmental from heritable effects. Real tumors often exhibit substantial subclonal mixing [34], bringing cells with different ancestries into close spatial proximity. Given a sufficiently large sample of cells, this mixing should allow us to fit a model that jointly accounts for spatial position and ancestry to distinguish their contributions to gene expression variation. Building on the joint modeling approach of Freckleton & Jetz, we develop Spatial Ornstein-Uhlenbeck Regression (SOUR) to accommodate gene expression counts and microenvironmental effects shared by cells regardless of their ancestry. In simulations with known ground truth, SOUR controls for the correlation between spatial and lineage distance when estimating the contribution of each component, whereas previously used methods exhibit substantial confounding. In two published spatial lineage tracing datasets, SOUR distinguishes spatial from heritable contributions, yielding more biologically plausible interpretations than approaches that analyze spatial and lineage information separately.

## 2 Results

### 2.1 Spatial Lineage Tracing Jointly Profiles Gene Expression, Location, and Ancestry

Once raw measurements have been appropriately pre-processed, spatial lineage tracing datasets contain three types of information. First, measurements of the expression of *p* genes in each of *n* cells, which we denote by the *n× p* matrix **Y**. Second, the spatial coordinates of the cells, which we denote by the *n× d* matrix **C**. Most experiments measure cells in two spatial dimensions, so typically *d* = 2. Third, ancestry information, which we denote by *T*, representing either clonal identities or a phylogenetic tree. The form of *T* depends on the lineage tracing technology. In *static* lineage tracing, clonal barcodes identify cells that shared a common ancestor at the time of barcoding [23, 26]. In *dynamic* lineage tracing, barcodes or naturally occurring genetic mutations evolve over time and allow phylogenetic trees to be reconstructed [21, 25]. SOUR treats each reconstructed tree as fixed and known.

To demonstrate the practical importance of modeling spatial position and ancestry jointly, we analyze two previously published spatial lineage tracing datasets generated *in vivo*. The first consists of five seqFISH [35] samples from four mice and contains WM989 cells, a *BRAF* V600E mutant human melanoma cell line [36], grown as subcutaneous xenografts in immunodeficient mice and statically lineage traced with the SpaceBar technology [23]. We refer to it as the *SpaceBar dataset*. The second consists of three MERFISH [37] samples from one mouse and contains 4T1 cells, a murine breast cancer model, grown as lung metastases following tail vein injection into an immunocompetent mouse and dynamically lineage traced with the PEtracer technology [21]. We refer to it as the *PEtracer dataset*.

### 2.2 Separate Analyses Can Misattribute Spatial Variation to Lineage and Heritable Variation to Space

Two approaches have previously been used to quantify spatial and lineage-associated variation in spatial lineage tracing datasets. Both analyze the association between gene expression and each source of variation separately rather than accounting for space and lineage jointly. Specifically, in the PEtracer dataset [21], separate autocorrelation statistics were calculated for each gene with respect to spatial position and the lineage tree (see Appendix A.9 for more details). And, in the SpaceBar dataset [23], each gene was assigned a *clone score* and a *space score*. The clone score quantified the range of average expression across clones, whereas the space score quantified the range of average expression across discrete spatial regions (see Appendix A.8 for more details). We refer to these approaches as the *marginal autocorrelation method* and the *marginal range method*, respectively. We refer to the clone score and space score as the *marginal clone range* and *marginal space range*, or simply the clone range and space range. We use *marginal* in its statistical sense: each method quantifies the association with one source of variation without conditioning on the other.

A central limitation of these approaches is that spatial proximity and cell relatedness are often strongly associated because sibling cells are born adjacent to one another. Published spatial lineage tracing datasets accordingly show substantial overlap between spatial and lineage neighborhoods (Fig. 1). As a result, gene expression variation that is entirely spatial can appear lineage-associated, while variation that is entirely lineage-associated can appear spatial.

**Fig. 1.**
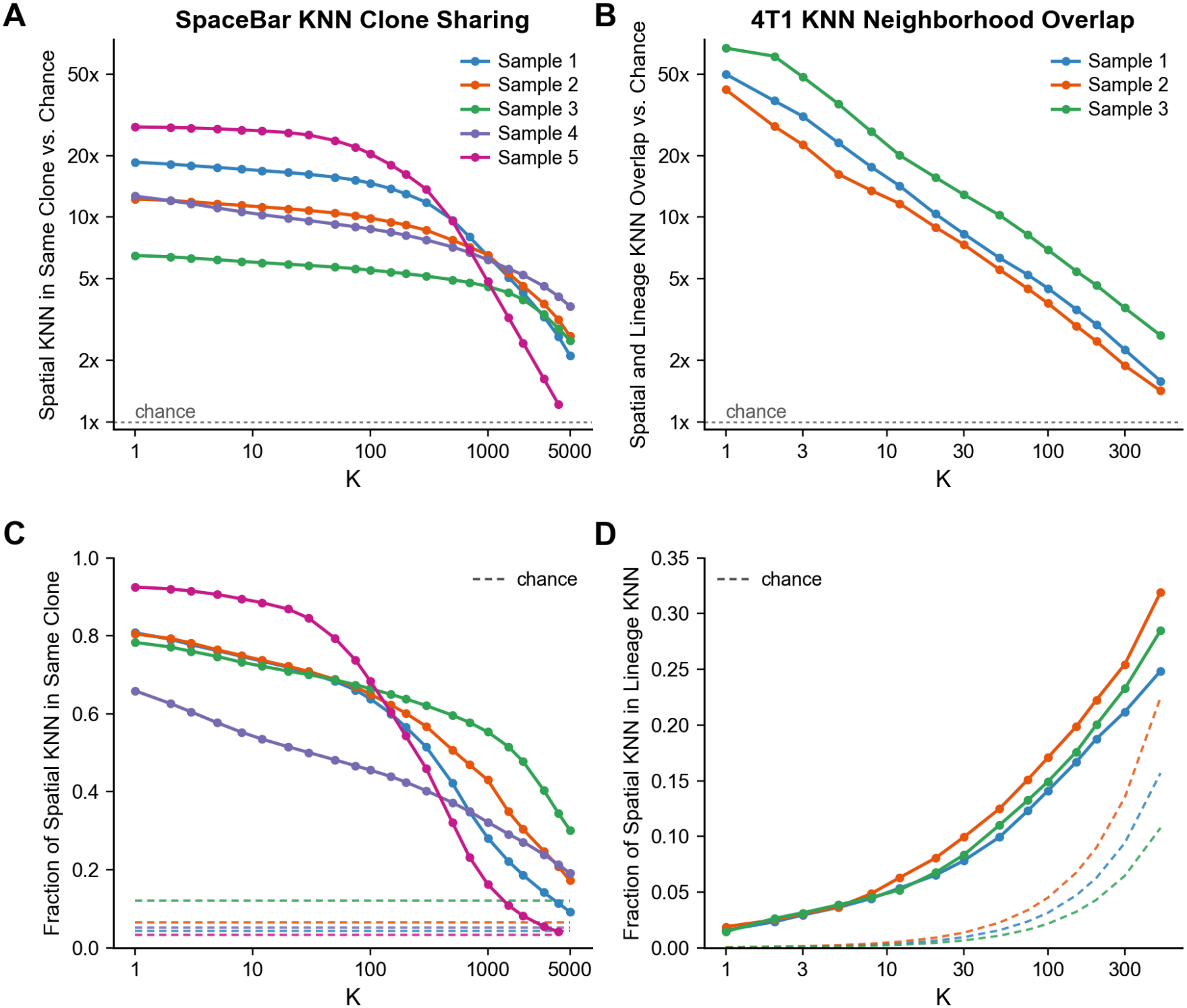
Lineage and spatial neighborhood overlap in two published datasets. Higher values indicate that spatially close cells are more related. **(A)** For each SpaceBar sample, the average proportion of a cell’s *K* nearest spatial neighbors that share the same clone as the given cell, normalized relative to the proportion expected by randomly sampling *K* other cells. **(B)** For each PEtracer sample, the average proportion of a cell’s *K* nearest spatial neighbors that are also among its *K* nearest lineage neighbors, normalized by the expected overlap of two randomly drawn sets of size *K*. **(C, D)** The same two statistics on the un-normalized proportion scale.

To test the extent to which separate analyses incorrectly attribute variation to space or lineage, we simulated data in which only one of these components contributed to gene expression heterogeneity. As described in Section 4.5, in the *spatial-only* simulation setting, we varied expression with the spatial positions of cells and with unstructured noise. And, in the *lineage-only* simulation setting, we varied expression with the lineage tree or clonal structure and with unstructured noise. Any variation attributed to lineage in the spatial-only setting, or to space in the lineage-only setting, therefore represents incorrect attribution.

To preserve the relationship between spatial position and ancestry found in real experiments, we retained the observed spatial coordinates and lineage information from the two published datasets mentioned above, and simulated only gene expression. Specifically, for sample 3 of the PEtracer dataset, we generated expression under the spatial-only and lineage-only settings and applied the marginal autocorrelation method. And, for sample 1 of the SpaceBar dataset, we generated expression under the same two settings and applied the marginal range method. In both simulation settings, we varied the strength of the contributing signal by manipulating the proportion of gene expression variation that it explained (as defined in Eq. 3). Further details of the simulation design are provided in Section 4.5.

Both methods attributed gene expression variation to the component that was absent (Fig. 2). As the contribution of the true component increased, the apparent contribution of the absent component also increased because spatial and lineage distances were correlated. Consequently, both the marginal range method and the marginal autocorrelation method produced high false-positive rates for lineage in the spatial-only simulations and for space in the lineage-only simulations (Fig. 2).

**Fig. 2.**
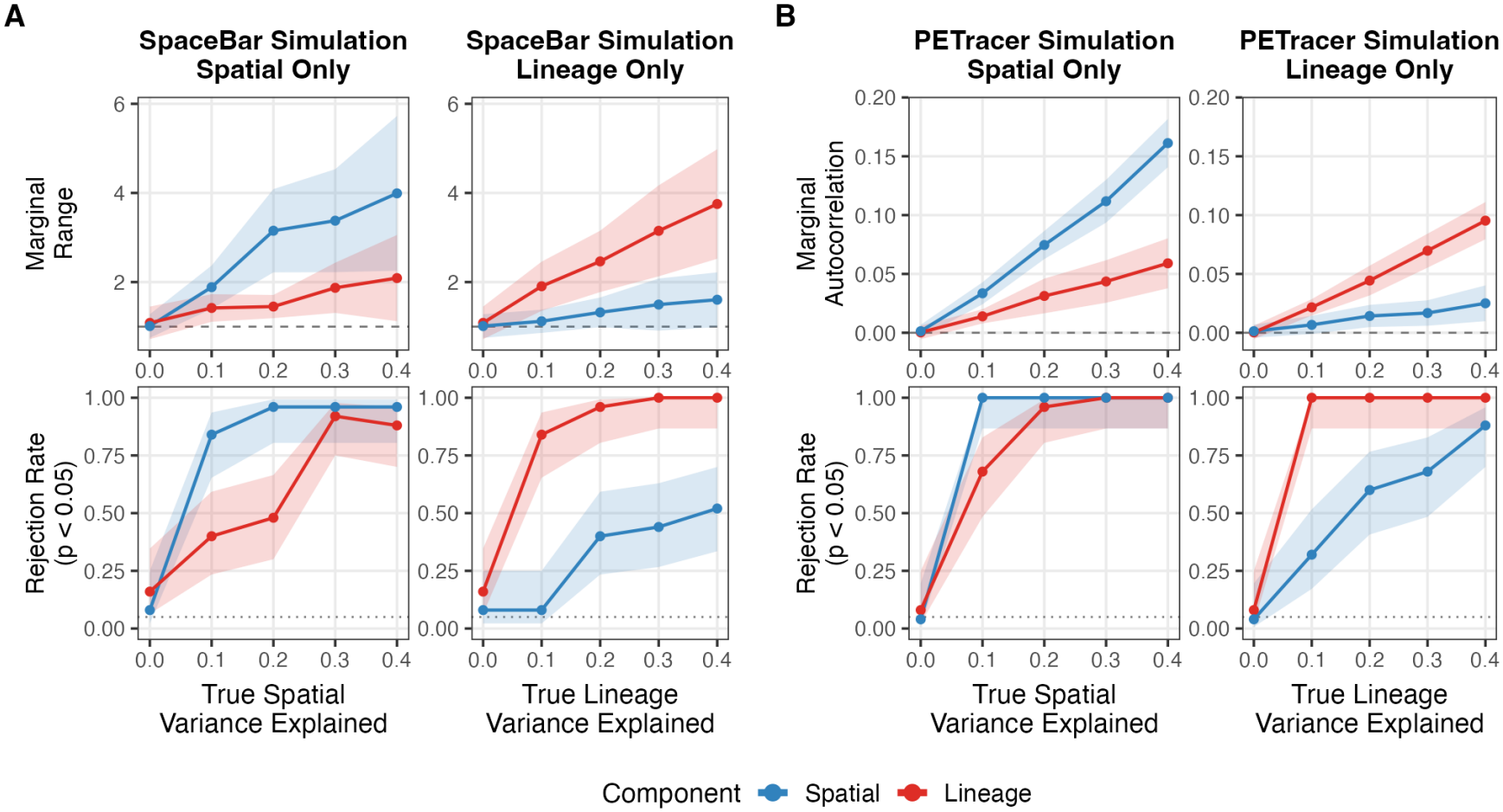
Results of applying previously used methods to simulated data across two settings using real spatial positions and lineage information from previously published datasets. The null hypothesis represented by the dashed line in the lineage-only simulations is that there is no relationship between lineage and gene expression, and the null hypothesis represented by the dashed line in the spatial-only setting is that there is no relationship between space and gene expression. In the bottom row, the dotted line marks the 0.05 level. **(A)** Results of applying the marginal range method to 25 replicate simulations using sample 1 of the SpaceBar data. In the top row, each point represents the average value, *±*1 standard deviation, of the test statistic of the marginal range method [23], where the null hypothesis implies an expected value of 1. The bottom row displays the average rejection rate at the 0.05 level of the p-value produced using the permutation test used in the marginal range method [23]. A 95% Wilson score interval is displayed for each point in the bottom row. **(B)** Results of applying the marginal autocorrelation method to 25 replicate simulations using sample 3 of the PEtracer data. The top row displays the average value, *±*1 standard deviation, of the autocorrelations, where the null hypothesis implies an expected value of 0, and the bottom row displays the rejection rate of a p-value (with associated 95% Wilson score interval) calculated by a permutation test.

Together, these simulations show that methods that analyze space and lineage separately can detect structured gene expression variation but cannot reliably identify which component accounts for it. This failure arises because the methods consider space and lineage in isolation even though the two are correlated. We therefore developed an approach that models gene expression variation as a function of space and lineage jointly.

### 2.3 SOUR Jointly Models Spatial and Lineage-Associated Gene Expression Variation

Multiple regression is the most straightforward approach for separating the associations of correlated predictors with an outcome. By including both predictors in one model, it estimates the association with each predictor while accounting for the other. Standard linear regression, however, requires predictors to be represented by numerical features and assumes a specified linear relationship between those features and the expected outcome.

Cell location and ancestry do not have natural representations that satisfy these requirements. Gene expression is unlikely to vary linearly across the spatial dimensions of a tissue. Similarly, it is unclear how to assign a numerical value to each cell that encodes its evolutionary relationship to other cells in a meaningful way. SOUR therefore represents space and lineage through covariance structures instead of treating them as ordinary regression covariates.

Specifically, SOUR decomposes latent gene expression variation across *n* cells into spatial variation, lineage-associated variation, and other cell-to-cell variation, while accounting separately for sampling variation in the observations. For a single gene, let **y** denote the vector of observed transcript counts. SOUR assumes that for each cell *i*,

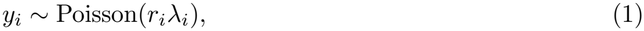

where *r*_*i*_ *>* 0 is a fixed scaling factor that accounts for differences in overall cell size or detection efficiency, and *λ*_*i*_ *>* 0 is the latent expression rate in cell *i*.

SOUR then decomposes the log expression rate for each cell *i* as

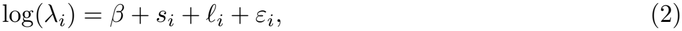

where *β* is an intercept, *s*_*i*_ is the spatial component of expression, *l*_*i*_ is the lineage-associated component of expression, and *ε*_*i*_ captures remaining cell-to-cell variation such as cell-cycle variation or inherent variability in gene expression [38]. To encode the assumption that nearby cells have correlated spatial components and closely related cells have correlated lineage components, we assume that the vector of spatial components **s** and the vector of lineage components ***l*** are drawn independently from Gaussian processes with Ornstein-Uhlenbeck covariances. In the special case when the experiment provides clonal identities rather than a full lineage tree, we instead assume that cells within the same clone share a lineage component, where the collection of lineage components across clones is drawn from a common normal distribution. The remaining cell-to-cell component is assumed to be independent and normally distributed across cells. We fit this model to the data **y** using a Bayesian framework, which provides a posterior distribution over ***l*, s**, and ***ε***. The full model, priors, identifiability constraints, and computational procedure are described in Section 4.

To compare the relative contributions of space, lineage, and other cell-to-cell variation, we first define theoretical summaries for the true latent components. These summaries exclude Poisson sampling variation in the observed counts, which is accounted for separately by the observation model. Let the vector ***η*** = **s** + ***l*** + ***ε*** denote the component of the latent log expression rate that varies across cells. Using Var_cell_ and Cov_cell_ to denote variance and covariance across the *n* observed cells, we define the *variance share* of each component as

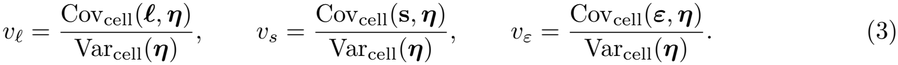

Each numerator measures how strongly one component varies with the total latent log expression rate. We use the covariance term in the numerator, as opposed to the variance of each component, to ensure that the three shares sum to one. The shares therefore allocate all latent variation across cells among the spatial, lineage-associated, and remaining cell-to-cell components. We interpret larger values of *v*_*l*_ to indicate that, accounting for spatial and unstructured variation, a larger proportion of ITH can be explained through the lineage component. *v*_*s*_ has an analogous interpretation.

In practice, the true latent components are unknown. To estimate Eq. 3, we therefore take the posterior distribution estimated by SOUR and then calculate the shares for each posterior sample of **s, *l***, and ***ε***. The resulting posterior distribution provides estimates of the relative contributions of the three components and quantifies uncertainty in those estimates.

By modeling spatial and lineage-associated variation jointly, SOUR is designed to separate the two sources when spatial proximity and cell relatedness are correlated. Separation becomes more difficult as the spatial and lineage structures become more similar, which increases uncertainty in the estimated variance shares. We therefore evaluate the accuracy, uncertainty, and robustness of these estimates in simulations with known contributions from each component in the following section.

### 2.4 Simulations Demonstrate that Current Sample Sizes Are Large Enough for SOUR to Accurately Estimate Variance Shares

Fitting the SOUR model to the same simulations used to create Fig. 2 demonstrates high accuracy of the estimated variance shares (Fig. 3). In contrast to both marginal methods, SOUR showed very little evidence of confounding. Specifically, the non-driving signal had average variance shares at or below 2.7% across all simulation scenarios. The variance shares of the driving signal and the noise component were also estimated quite accurately, showing only very slight negative biases for large spatial shares and very slight positive biases for the PEtracer lineage-only simulation. Overall, this simulation study suggests that the number of cells measured in these real datasets is large enough for SOUR to overcome the lineage-spatial correlation structures demonstrated in Fig. 1.

**Fig. 3.**
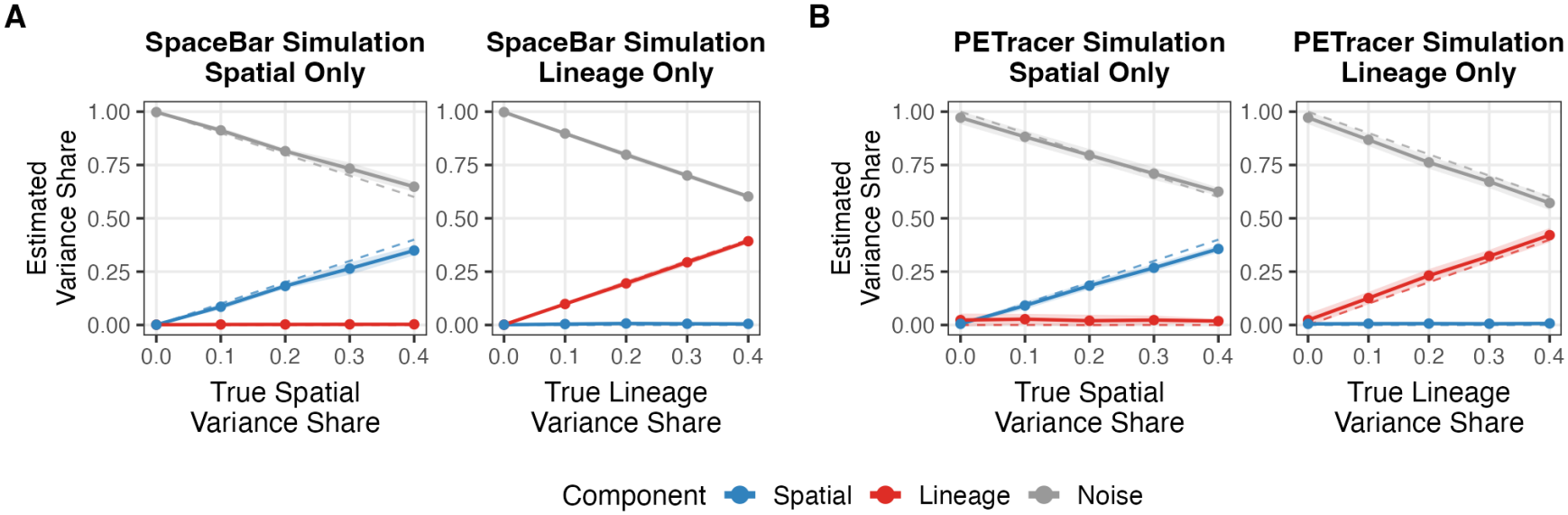
Results of fitting SOUR on the same simulated data as Fig. 2. Each point is the average over replicate simulations of the posterior mean variance share, and error bands are *±*1 standard deviation over the replicates. Dashed lines represent true variance shares. **(A)** Simulation results on the SpaceBar dataset. **(B)** Simulation results on the PEtracer dataset.

We note that while our simulations used the Poisson likelihood (Eq. 1) and additive decomposition (Eq. 2) of the SOUR model, we generated the SpaceBar spatial-only data and PEtracer lineage-only data from a generative process that *differed* from the exact Gaussian process kernels that the SOUR model assumes (we describe these kernels, and how our simulations differed from them, in Section 4.2 and Section 4.5, respectively). Thus, these simulations demonstrate that SOUR does not require all of its modeling assumptions to be met in order to return accurate variance shares.

In Appendix B.1, we provide extended results from the simulation study presented in Fig. 3, as well as additional simulation studies investigating other forms of model misspecification. These simulations further demonstrate that SOUR avoids the confounding of previous methods across a wide array of scenarios, even when parts of the SOUR model are misspecified. In some cases, however, SOUR can be slightly conservative, attributing some of the structured variance to unstructured noise.

### 2.5 Reassessment of Spatial and Lineage Contributions to Heterogeneity in Published Tumor Datasets with SOUR

#### 2.5.1 SOUR Clarifies the Role of Immediate Early Genes in a Melanoma Experiment

Here, we re-analyzed the 5 tissue samples of the SpaceBar dataset [23]. We fit SOUR separately to each gene in each sample, as opposed to fitting to each gene in all samples simultaneously, because the samples contain disjoint clonal barcodes sampled from different tumors. We analyzed all genes detected in at least 5% of a sample’s cells, which left between 90 and 108 genes per sample.

Overall, SOUR attributed substantially more gene expression variation to spatial structure than to clonal identity in this system. Across genes and samples, the average spatial variance share was 31.7% (95% CI 29.1% to 34.3%), compared with an average clonal variance share of 6.5% (95% CI 5.6% to 7.3%). The remaining variation was attributed to other cell-to-cell differences (Table 1). This broad conclusion agrees with the original SpaceBar analysis [23]: in four of the five samples, the mean space range exceeded the mean clone range, and a larger proportion of genes was classified as spatially significant than clonally significant (Table 1).

**Table 1.**
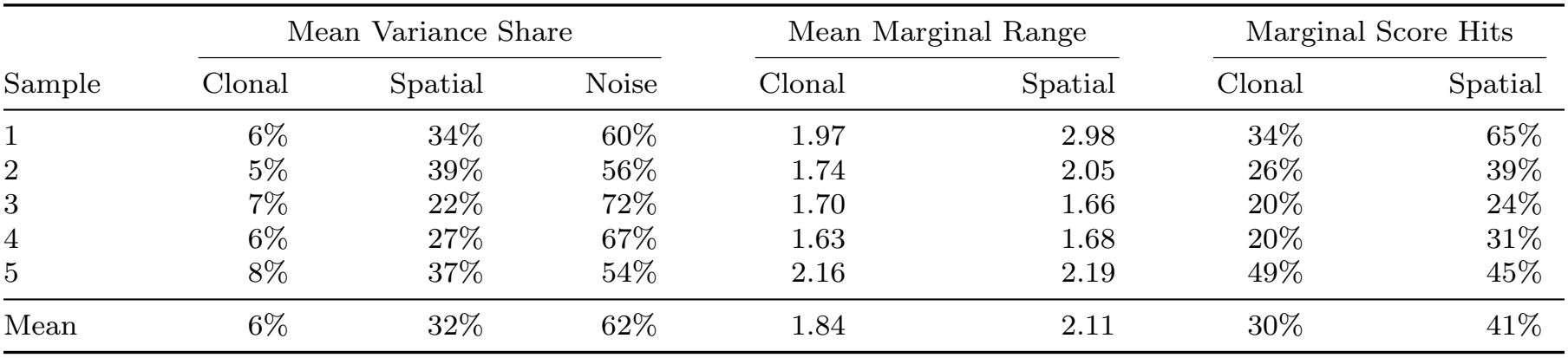
Posterior mean variance shares, as percentages, in each sample of the SpaceBar dataset, averaged over the genes analyzed in each sample, together with the mean SpaceBar marginal ranges over the same genes and the percentage of those genes that the original publication called significant (i.e., Marginal Score Hits).

Although both methods attributed more variation to space than to clonal identity when results were aggregated across genes, their gene-level results showed little agreement, especially in sample 3 (Fig. 4A,B; Fig. C10). Across the 108 genes, the Pearson correlation between SOUR’s spatial variance shares and the space ranges was *ρ* = 0.20 (95% CI 0.01 to 0.37), and the corresponding correlation for the clonal measures was *ρ* = 0.00 (95% CI *−*0.14 to 0.17). One possible reason for this discrepancy is that SOUR calculates an *average* contribution over the entire sample, whereas the marginal range method is only a function of the *extreme* clones and spatial bins. To test this explanation, we also compared the marginal range method to the posterior range of the fitted SOUR model, defined as E[max_*i*_ *s*_*i*_ *−* min_*i*_ *s*_*i*_ |**y**] for the spatial field and analogously for the clonal effects. Using this posterior summary, as opposed to the variance shares, did not resolve the broad disagreement between the methods (Fig. C11), suggesting that the discrepancy is a result of fundamental differences in the modeling approaches as opposed to the choice of posterior summary.

**Fig. 4.**
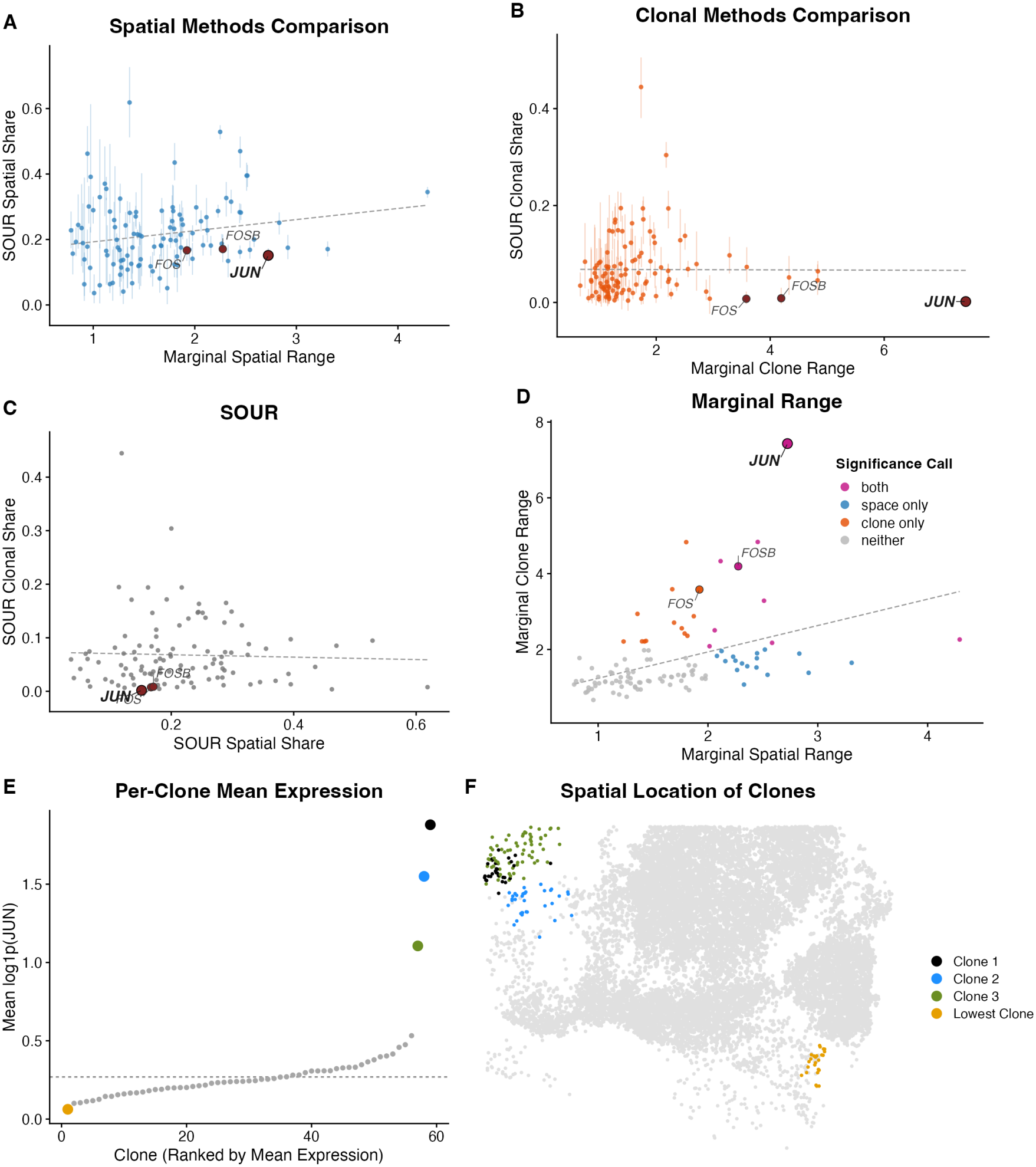
Results of SOUR and the marginal range method on sample 3 of the SpaceBar dataset. **(A, B)** SOUR posterior mean spatial and clonal variance shares (with associated 95% credible intervals) against the corresponding clone and space ranges originally published in [23]. Dashed lines are least squares fits. **(C, D)** Clonal and spatial measures for each method plotted against each other over the same genes, SOUR variance shares in (C) and marginal ranges in (D). Points in (D) are colored by the significance calls made in the original study. Dashed lines are least squares fits. **(E)** Mean normalized *JUN* expression, log(1 + *y*_*i*_*/r*_*i*_) with *r*_*i*_ the total counts in cell *i* scaled to a median of 1 across cells, within each of the 59 clones, ranked, with the mean across all cells shown as a dashed line. **(F)** Spatial positions of cells in sample 3 of the SpaceBar dataset, with the three highest-expressing clones and the lowest-expressing clone colored as in (E).

The largest single-gene discrepancy in sample 3 illustrates how SOUR separates spatial from clonal variation: *JUN* ranked first by clone range but last by posterior mean clonal variance share (approximately 0.2%), because its apparent clonal variation is better explained by space. The mean normalized expression (*y*_*i*_*/r*_*i*_) of *JUN* in the top-ranked clone was 72*×* that of the lowest-ranked clone (30 *×* on the log(1 + *y*_*i*_*/r*_*i*_) scale of Fig. 4E), so the clone range was very large. However, the three highest-ranked clones occupy a distinct spatial neighborhood (Fig. 4F), and SOUR accordingly attributed this variation to space, fitting very high values of the spatial field **s** in this neighborhood (Fig. C12A). This attribution appears to be well justified given the experimental design. Because clones are defined by independently assigned barcodes in a common parental population, there is no reason to expect neighboring clones to be more related than distant ones. Three adjacent clones independently experiencing genetic or epigenetic events that up-regulate *JUN* is therefore far less plausible than a shared local spatial effect.

Functionally, our analysis above suggests that there is a spatially restricted microenvironmental force heavily up-regulating the expression of *JUN*. Notably, we found that the tissue region that showed up-regulation of *JUN* also showed marked up-regulation of both *FOS* and *FOSB* (Fig. C12). Together, *JUN, FOS*, and *FOSB* are classified as *immediate early genes* (IEGs) [32], which are known to be rapidly responsive to external signaling cues. IEG transcription is typically initiated by signals a cell receives from its surroundings, such as growth factors or the presence of oxidative stress, rather than by cell-intrinsic programs [32, 33]. Thus, it is also biologically more plausible that this IEG up-regulation is a result of a microenvironmental force as opposed to an inherited trait.

Finally, we note that across genes, the SOUR spatial and clonal variance shares were far less correlated than the space and clone ranges (*ρ* = *−*0.04, 95% CI *−*0.18 to 0.14, vs. *ρ* = 0.42, 95% CI 0.32 to 0.54; difference in absolute correlation *−*0.38, 95% CI *−*0.50 to 0.19; Fig. 4C,D). Given that the SOUR spatial variance shares are much larger than the SOUR clonal variance shares, many of the previously published clone ranges may mostly be a result of confounding.

#### 2.5.2 SOUR Attributes *Slc2a1* Variation to Spatial Hypoxia Rather Than Lineage

We also applied SOUR to three samples from a single mouse in the PEtracer dataset. We analyzed 39 of the 175 measured genes, filtering out genes we suspect were frequently misassigned to the tumor cells as a result of segmentation error. We describe our filtering criteria in Appendix A.7. The three samples share a lineage tree but lie in different coordinate planes, so we allowed each sample to have a separate spatial field, with shared hyper-parameters across these fields, while retaining a shared lineage structure, as described in Appendix A.6.

Overall, the marginal autocorrelation method suggests that space and lineage are similarly important in determining heterogeneity in this system, whereas the SOUR method strongly favors space. Across the 39 genes, the average spatial variance share was 14.5% (95% CI 12.3% to 16.7%) whereas the average lineage variance share was 4.9% (95% CI 4.0% to 5.8%), while the average spatial autocorrelation was 0.085 (95% CI 0.072 to 0.099) and the average lineage autocorrelation was 0.079 (95% CI 0.059 to 0.098) (Fig. 5A,B). Across all 39 genes, the two marginal autocorrelations were themselves correlated: *ρ* = 0.47 (95% CI 0.31 to 0.88), rising to *ρ* = 0.74 (95% CI 0.48 to 0.90) when excluding *Cldn4*, the gene with by far the greatest lineage autocorrelation (Fig. 5D). This is the pattern confounding would produce if space were the driving signal: because related cells lie close together, a gene with strong spatial structure will also register a high lineage autocorrelation. In contrast, the SOUR lineage and spatial variance shares were only weakly correlated across genes, *ρ* = 0.16 (95% CI *−*0.16 to 0.50) (Fig. 5C).

**Fig. 5.**
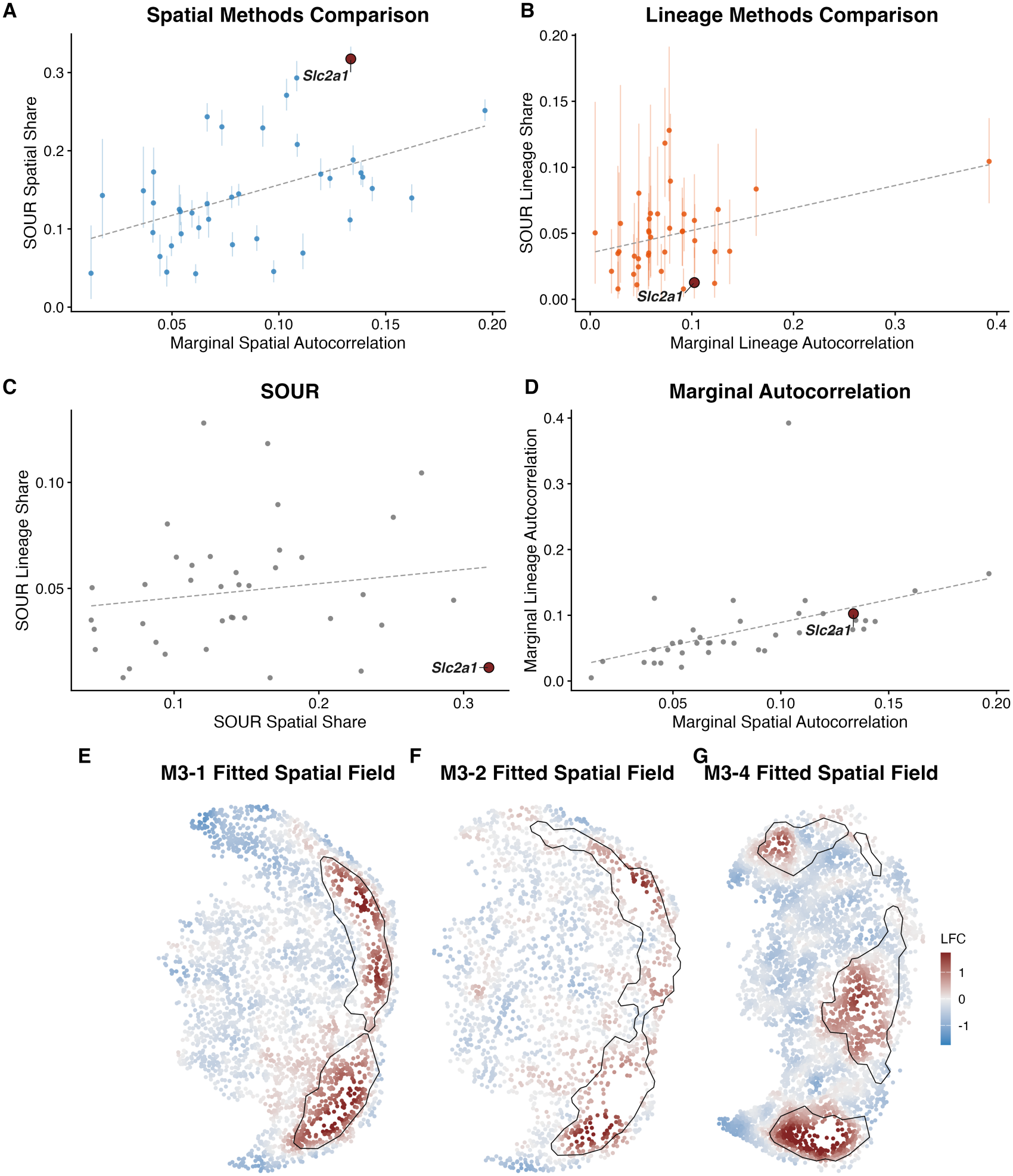
Results of SOUR and the marginal autocorrelation method on the PEtracer dataset. **(A, B)** SOUR posterior mean spatial and lineage variance shares (with associated 95% credible intervals) against the corresponding marginal autocorrelations originally published in [21]. Dashed lines are least squares fits. **(C, D)** Lineage and spatial measures for each method plotted against each other over the same genes, SOUR variance shares in (C) and marginal autocorrelations in (D). Dashed lines are least squares fits. **(E–G)** Posterior mean of the spatial field **s** of the SOUR model fit to *Slc2a1*. The outlined region is the hypoxia module originally identified and shown in Figure 6C of [21]. Values represent a log-fold change relative to the expected expression due to the intercept *β* and lineage component ***l***.

The two methods sometimes produced very different results for individual genes (Fig. 5A,B; Fig. C13). While the two methods showed moderate positive correlation across genes (lineage measures *ρ* = 0.36, 95% CI 0.02 to 0.63; spatial measures *ρ* = 0.47, 95% CI *−*0.24 to 0.65), large differences in the results on individual genes remained. For example, the gene *Slc2a1* had relatively similar values of spatial and lineage autocorrelation (0.134 and 0.102, respectively), whereas SOUR strongly favored the spatial variable, attributing 31.7% of the variance to space and only 1.3% of the variance to lineage, with the rest being attributed to noise. Interestingly, we found that high values of the fitted spatial component for *Slc2a1* overlapped strongly with the hypoxia module identified in the original PEtracer analysis across all three samples (Fig. 5E–G) [21]. This correspondence is biologically consistent with regulation of *Slc2a1* by hypoxia-inducible factor 1*α* (*Hif1a*) [39]. Cells assigned to the hypoxia module showed some local clustering on the lineage tree but were distributed across multiple parts of the tree (Fig. C14). Together, the spatial expression pattern, overlap with the independently defined hypoxia module, and broad distribution across the lineage tree support a predominantly spatial interpretation of *Slc2a1* variation in this experiment.

## 3 Discussion

Spatial lineage tracing creates an opportunity to distinguish gene expression variation associated with the local environment from variation associated with shared ancestry. It also creates a statistical challenge because related cells tend to remain spatially close. We developed SOUR to address this dependence by modeling spatial position and ancestry jointly and partitioning latent gene expression variation into spatial, lineage-associated, and other cell-to-cell components. In simulations, approaches that analyzed space and lineage separately attributed signal to components that were absent, whereas SOUR more accurately recovered the contributions used to generate the data. These results establish that joint modeling is necessary when spatial and lineage relationships are correlated and show that SOUR can separate their contributions in many cases when the data contain sufficient information to do so.

The analyses of two published datasets illustrate how joint modeling changes biological interpretation. In the SpaceBar melanoma experiment, the clone range ranked *JUN* as strongly clone-associated, whereas SOUR assigned it little clonal variation and identified a localized spatial pattern shared by *JUN, FOS*, and *FOSB*. The coordinated spatial expression of these immediate early genes is consistent with a local extracellular stimulus. In the PEtracer experiment, SOUR attributed *Slc2a1* variation predominantly to space rather than lineage. Its fitted spatial component overlapped a hypoxia module identified independently in the original analysis, consistent with the known regulation of *Slc2a1* by *Hif1a*. These examples show how analyzing either structure alone can produce a plausible but potentially misleading biological interpretation.

Notably, in both datasets SOUR attributed several times more gene expression variation to spatial position than to ancestry, suggesting that in these systems ITH is shaped more by the microenvironment than by heritable differences between cells. Whether the same holds in human tumors (or even in other model systems) is unclear. Both experiments followed tumors grown in mice from established cell lines for four to six weeks, whereas human tumors accumulate heritable genetic and epigenetic diversity over years of growth and selection [4, 40, 41].

The components estimated by SOUR describe statistical associations rather than mechanisms. A spatial component captures expression variation organized by physical location, but it does not by itself identify a microenvironmental cause. Such variation could reflect the spatial distribution of stromal or immune cell types, gradients in oxygen or nutrients, or local signaling. Similarly, a lineage-associated component need not reflect a genetic change; it can arise from any persistent state shared among related cells. Experimental perturbations or additional molecular measurements are required to identify the processes responsible for either component. SOUR is therefore best viewed as a method for identifying genes and spatial regions for mechanistic follow-up.

SOUR also complements a growing set of methods for spatial lineage tracing data. Some previous methods, such as MOLLUSC, BEAM, and LineageMap, focus primarily on lineage-tree reconstruction and estimating spatial trajectories of cells [42–44]. While LineageMap can additionally model the evolution of cell states, it requires clustering cells into discrete types before analysis, which may be a poor fit for cancer datasets that typically exhibit a spectrum of states. SOUR instead models gene expression directly and is designed to estimate the distinct contributions of spatial and lineage structure to a continuous molecular phenotype.

The SOUR framework also creates several opportunities for future development. The current implementation conditions on reconstructed lineage trees, clone assignments, and cell segmentation, allowing it to be applied directly to the outputs of existing spatial lineage tracing pipelines. Future extensions could propagate uncertainty in barcode detection, tree reconstruction, and cell segmentation through to the estimated variance shares. In addition, SOUR currently fits each gene independently, ignoring coordinated expression between genes. Multivariate or hierarchical extensions of SOUR could share information across co-regulated genes, improve estimation for sparsely measured transcripts, and identify spatial or heritable expression programs directly. Finally, while the additive structure of SOUR provides an interpretable framework for decomposing spatial, lineage-associated, and remaining cell-to-cell effects on the latent log expression scale, it ignores potential interactions between these factors. For instance, hypoxia can induce genetic instability [45–47] and persistent epigenetic changes [48]. Extending SOUR to represent interactions could identify genes for which environmental responses depend on ancestry. The current decomposition supplies both a benchmark for these extensions and a way to determine whether the available data support the additional complexity.

## 4 Methods

### 4.1 Notation

Below, we use bolded capitalized letters to indicate matrices (e.g., **X**), bolded lowercase letters to indicate vectors (e.g., **x**), and unbolded lowercase letters to indicate scalars (e.g., *x*). We define our model below with the notation introduced in Section 2.1.

### 4.2 Statistical Model

For a single gene, SOUR decomposes variability in the latent log rates that produced the gene expression counts **y** into three components: a lineage term ***l***, a spatial term **s**, and the term ***ε***, accounting for other unexplained variability and assumed to be independent and identically distributed (iid).

We formalize this decomposition with the following generalized linear mixed effects model (GLMM):

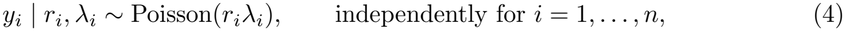

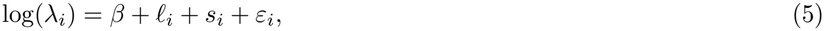

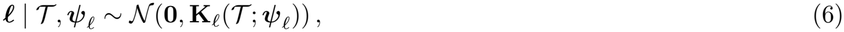

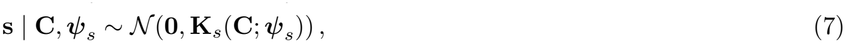

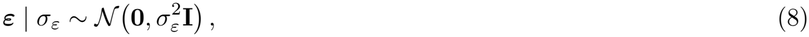

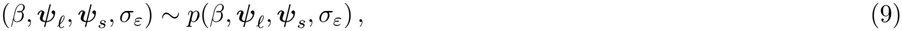

where *r*_*i*_ *>* 0 is a fixed scaling factor that accounts for cell size, *β* is an intercept, and **K**_*s*_ and **K**_*l*_ are covariance matrices that are functions of known covariates (**C** and *T*, respectively) and hyperparameters (***ψ***_*s*_ and ***ψ***_*l*_, respectively). In our experiments, we define *r*_*i*_ to be the total counts across all genes in cell *i* normalized to have a median of 1 across cells. The intercept *β* and hyper-parameters ***ψ***_*l*_, ***ψ***_*s*_, and *σ*_*ε*_ are drawn from a fixed prior distribution as described in Appendix A.3.

Below, we describe our parameterizations of the covariance matrices **K**_*l*_ and **K**_*s*_. We note that we use a different form for the lineage covariance depending on whether the experiment allows for the reconstruction of full phylogenetic trees or merely clonal barcodes.

#### 4.2.1 Spatial Covariance

We parameterize the spatial covariance **K**_*s*_ with 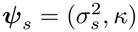, where 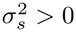 represents the marginal spatial variance and *κ >* 0 represents the inverse spatial length-scale. Applying the exponential kernel (also known as the Ornstein-Uhlenbeck or OU kernel) to the matrix of spatial positions **C** [49, Section 4.2], we obtain the unconstrained covariance matrix 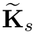, defined entry-wise as

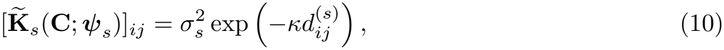

where 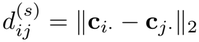 is the Euclidean distance between cells.

As described in Appendix A.1, using 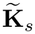 in Eq. 7 could cause an identifiability issue between **s** and *β* in certain regions of the parameter space. To avoid this issue, we impose the constraint that **s** sums to 0, which is achieved by defining

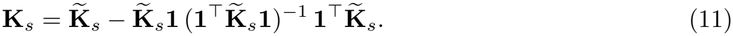

#### 4.2.2 Lineage Covariance – Phylogenetic Tree Case

In the case where *T* consists of a rooted, phylogenetic tree, we derive the lineage covariance of the observed cells **K**_*l*_ from assuming that the latent lineage values along the tree follow a phylogenetic OU process. Such processes have been used extensively to model the evolution of traits among related species [50], and more recently to model the evolution of gene expression over lineage trees [51, 52]. At a high level, the phylogenetic OU process assumes that the latent lineage component of expression evolves along the tree according to a stochastic process that involves both random variation as well as a restoring force towards a fixed value. This process occurs in continuous time, but in our model the values *l*_1_, …, *l*_*n*_ represent the states of the OU process of cells observed at the time of measurement, *t*_obs_. We describe the phylogenetic OU process in more detail in Appendix A.2.

Here, we parameterize the OU process with 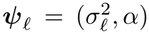, where 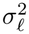 represents the marginal variance of the lineage component and *α >* 0 represents the restoring force. Under the assumption that the latent lineage component of the ancestral root cell at time *t* = 0 is fixed at 0, we can write the unconstrained covariance matrix of the lineage values at the leaves of the tree, 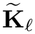, entry-wise as

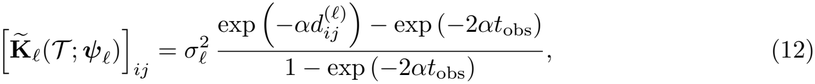

with 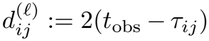, where *τ*_*ij*_ denotes the time from the root of *T* to the most recent common ancestor of cells *i* and *j*. We justify and further discuss our parameterization of the OU process and the assumption of the fixed root in Appendix A.2. We note that in Eq. 12, the functional form of the first term of the numerator is the same as the unconstrained spatial covariance (Eq. 10), and in the limit where *t*_obs_ *→ ∞*, the two covariances have the same form.

Finally, for consistency with the spatial covariance, and because 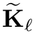 could cause identifiability issues between *β* and ***l***, we define

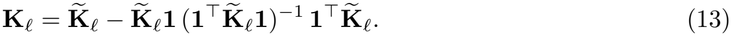

#### 4.2.3 Lineage Covariance – Clonal Case

When a spatial lineage tracing experiment was performed with static barcodes, such as in [23, 24, 26], *T* will not contain a full rooted phylogenetic tree, but rather only information about which cells share a common ancestor. Specifically, our population of cells will be marked at some point in time with one of *B* heritable barcodes, which we label 1, …, *B*. Then, at the end of the experiment, for each cell *i* we read out a barcode *b*_*i*_ *∈ {*1, …, *B}* .

To capture inherited clonal variation, we assign each *clone* a latent effect

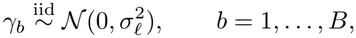

and set 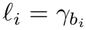, so that cells carrying the same barcode share a lineage component. Therefore, for any two cells *i* and *j*, the covariance is parameterized by 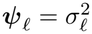, and we have

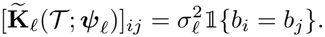

To ensure identifiability with the intercept *β*, we again require that the sum of the values of ***l*** is equal to 0. Writing 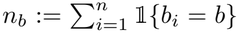 for the number of cells carrying barcode *b*, we define the constrained covariance as

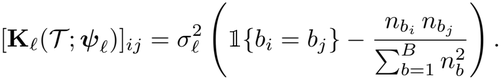

We note that there is unfortunately no direct correspondence between the clonal covariance and the phylogenetic OU covariance. In general, we expect that using clonal barcodes in an experiment will result in smaller estimates of the lineage variance shares than had the same experiment been run with evolving barcodes that permit tree reconstruction, as all intra-clonal variation will be lost and is likely to look like unstructured noise.

### 4.3 Posterior Inference

Given the observed data **y**, we would like to calculate the following posterior over the model parameters:

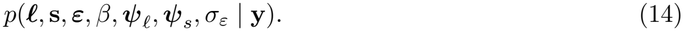

Unfortunately, direct computation of Eq. 14 is intractable, so we instead use INLA [53] to approximate this distribution. While alternative MCMC approaches would eventually become arbitrarily accurate if given enough running time, INLA will suffer some approximation error, no matter how much running time it is given, due to its underlying assumptions. However, INLA is much faster than MCMC, and is particularly well suited for providing accurate approximations to the class of GLMMs to which SOUR belongs. We discuss further details of this approach in Appendix A.4.

### 4.4 Model Interpretation

As described in Section 2.3, we summarize the results of the fitted SOUR model by computing posterior quantiles of the variance shares defined in Eq. 3. Formally, defining

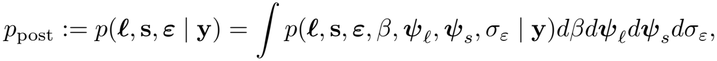

we compute

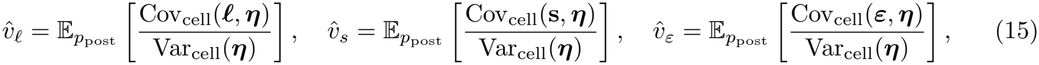

as well as posterior quantiles of the variance shares in our results. Here, Var_cell_ and Cov_cell_ are variances and covariances over cells, respectively, defined for *n*-vectors **u** and **w** as:

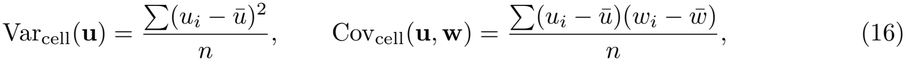

where 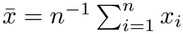. Note that because of the linearity of expectation, 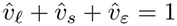.

In practice, we compute Monte Carlo estimates to Eq. 15 and other posterior quantiles of the variance shares using samples drawn from the approximate posterior obtained by fitting the model with INLA.

### 4.5 Generation of Simulated Data

In each of our simulations, we first generate the latent variables ***l*, s**, and ***ε***, and then generate observed gene expression counts **y** from the Poisson observation model (Eq. 4). We always set the value of the intercept, *β*, to 0, and we set the values of the scaling constants, *r*_*i*_, to be the total counts observed in each cell in the real datasets we use for simulation, normalized to have a median of 1.

Our simulation scenarios differ in exactly how the latent variables are generated. In each scenario, we target some pre-determined variance shares 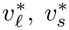 and 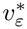 (where 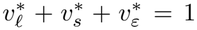). Conveniently, because the variance shares are a function of the empirical variances and covariances of the realized values of the latent variables, we can first generate candidate values of the latent variables from any non-atomic distribution such that ***l*** ⫫**s** ⫫ ***ε***, and then normalize these values to have (approximately) the desired variance shares.

Concretely, suppose that we have drawn candidate values of the latent variables 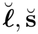, and 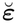. Then, for any positive value of 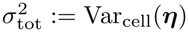, we can set

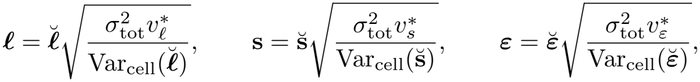

which implies

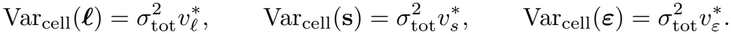

Recalling that the variance share for a given component **u** is defined as

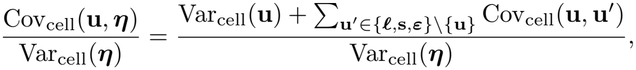

the above simulation setup results in

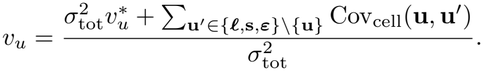

For a given realization of ***l*, s**, and ***ε***, it will not be the case that the covariances Cov_cell_(***l*, s**), Cov_cell_(**s, *ε***), and Cov_cell_(***l, ε***) are exactly 0. However, because ***l*, s**, and ***ε*** are generated independently, this will approximately be the case, and thus 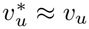 for **u** *∈ {****l*, s, *ε****}*. In all of our simulations, we set 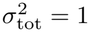.

In our simulations, we generated 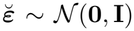. In the lineage-only simulations shown in Fig. 2 and Fig. 3, we set 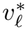 to the value on the x-axis and 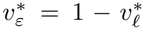, with 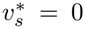. We set these values analogously for the spatial-only simulations. We describe how we generated 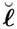and 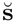for these simulations in Appendix A.5.

## Supporting information

Supplemental Data 1: Variance Shares

## Acknowledgements

The work of E.W. was funded in part by an MIT Health and Life Sciences (HEALS) Graduate Fellowship. The work of K.J.A. was funded in part by the NIH Pediatric Scientist Development Program (PSDP). The work of R.A.I. was funded in part by NIH grant R35GM131802.

## Data availability

The PEtracer dataset was obtained using the function datasets.koblan25() in the pycea package in python [54]. The SpaceBar dataset was obtained upon request from the authors of the original manuscript. The per-gene variance decompositions estimated by SOUR for each sample in both datasets are provided as Supplementary Data 1.

## Code availability

SOUR is implemented in the R package sour, available at https://gitlab.com/ericweine/sour together with installation instructions. Scripts reproducing all analyses, simulations and figures in this paper are available at https://gitlab.com/ericweine/sc_lineage_modelling.

## Author contributions

E.W. conceived the project. E.W., D.B., R.A.I., T.B., and J.S. developed the methods. E.W. implemented the methods and generated the figures. E.W., D.B., K.J.A., W.C., J.W., R.A.I., T.B., and J.S. wrote the paper.

## Appendix A Supplementary Methods

### A.1 Sum-to-Zero Identifiability Constraints

In Section 4.2, we describe sum-to-zero constraints imposed on the spatial term **s** and lineage term ***l***. Our primary motivation for applying these constraints is that, in their absence, **s** and ***l*** can become difficult to distinguish from the intercept *β*. Taking the spatial field as an example, the vector of rates ***λ***, and thus the likelihood of the data **y**, is unchanged under the transformation (*β*, **s**) *→* (*β −δ*, **s** + *δ***1**) for any *δ* ∈ ℝ. The likelihood therefore determines only the sum 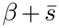, where 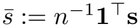 is the average spatial effect, and the division of this sum between *β* and 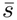 is determined by the prior. Because the prior on *β* is essentially flat, the prior on 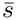 is primarily responsible for determining this division. Under the unconstrained spatial covariance (Eq. 10), 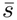is normal with mean 0 and variance

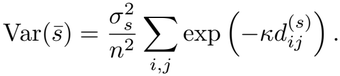

When the length-scale 1*/κ* is short relative to the extent of the tissue, the off-diagonal terms of this sum are small and 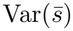 is close to 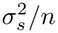 . The prior then holds 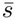 near 0 and assigns any level shared across all cells to *β*. However, when *κ* 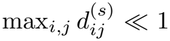, every term in the sum is close to 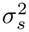 and 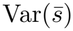 is close to 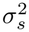 A draw of **s** from the prior is then approximately the constant vector 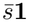 with 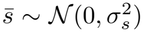 . Thus, in regions of the hyper-parameter space where *κ* is sufficiently small and *σ*_*s*_ sufficiently large, neither the likelihood nor the prior distinguishes between a shared level carried by *β* and one carried by **s**.

Identification of both **s** and *β* is primarily important when directly inspecting the posterior moments of **s**, as we do for the posterior mean of **s** in Fig. 5. When using the unconstrained covariance in the spatial prior (Eq. 10), it is possible that the posterior mean of **s** could be nearly uniformly large (either positive or negative), which would lead to the counterintuitive conclusion that the expression of a gene is being up-regulated or down-regulated across the entire tissue. By instead using the constrained spatial covariance (Eq. 11) that forces 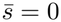, each value *s*_*i*_ can be interpreted as spatial regulation *relative* to the average cell. A similar argument can be made for the sum-to-zero constraint on the lineage term ***l***.

We note that the sum-to-zero identifiability constraints should not influence the variance shares under the posterior (Eq. 15), since the mean of each component is subtracted when calculating its variance and covariances across cells (Eq. 16).

### A.2 Fixed-Root OU

Here, we give a full description of the process and assumptions that result in the unconstrained phylogenetic lineage covariance (Eq. 12).

Recall that *T* contains a full, rooted phylogenetic tree with *n* leaves corresponding to the *n* observed cells. To aid in notation, we write *T* = (*V, E* ), where *V* denotes the set of vertices, including both internal nodes and leaves, and *E* denotes the set of directed edges. Let *a*_0_ ∈ *V* denote the root vertex and let *L* ⊂ *V* denote the set of leaves, with |*L*| = *n*. Similarly, let *I* := *V \* (*L* ∪ *{a*_0_*}*) denote the set of internal, non-root nodes. For each observed cell *i*, let *v*_*i*_ ∈ *L* denote the leaf corresponding to that cell. We write the ancestral vertices as *a*_0_, *a*_1_, …, *a*_|*I*|_, ordered by their split times.

We assume that the tree is *ultrametric*, meaning that all observed cells are sampled at the same evolutionary time. We assume that the root splits at time *t* = 0 and all leaves are observed at time *t* = *t*_obs_. Thus, each vertex *v* ∈ *V* has an associated birth time *t*_*v*_ ∈ [0, *t*_obs_], with 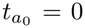 for the root and 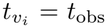 for all leaves *v*_*i*_ ∈ *L* . For a directed edge (*p, c*) ∈ *ε*, where *p* is the parent of *c*, the corresponding branch represents the time interval

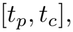

and has branch length

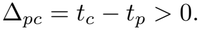

Internal vertices correspond to cell division events where a given node *v* splits into 2 children (in practice, the reconstructed tree *T* may include “polytomies”, or splits in the tree with more than 2 children; our model handles these without modification).

We model the lineage component of expression as a latent stochastic process that evolves forward in time along the branches of *T* . While this process takes place in continuous time, for convenience we work with its values at the vertices of the tree. Let

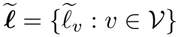

denote the collection of latent lineage states at all vertices. The lineage effects appearing in Eq. 6 are then the leaf states,

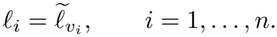

Fig. A1 illustrates this notation on a small tree with four leaves.

**Fig. A1.**
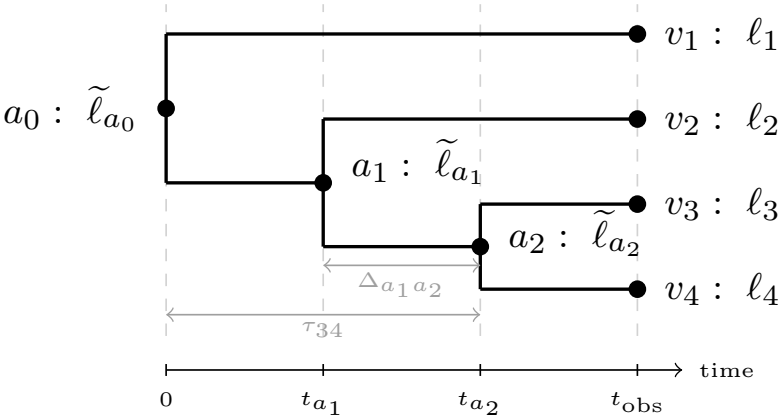
Tree notation on an example with *n* = 4 leaves. Time runs left to right, so the horizontal extent of a branch is its length in time: the root *a*_0_ splits at *t* = 0 and every leaf is observed at *t*_obs_, making the tree ultrametric. Filled circles are vertices *V*, with ancestral vertices *a*_1_, *a*_2_ ∈ *I* and leaves *v*_1_, …, *v*_4_ ∈ *L*. Each vertex *v* carries a latent state 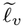, and the lineage effects entering Eq. 6 are the leaf states 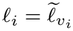 . The branch (*a*_1_, *a*_2_) has length 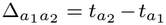 . Cells 3 and 4 have most recent common ancestor *a*_2_, so 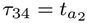 and their tree distance is 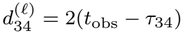.

We assume the stochastic process over 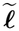 starts with a root state, 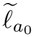, which we assume is fixed (see below for further discussion of this choice). At time *t* = 0, this root cell splits into two cells, *c*_1_ and *c*_2_. For the lifetime of cell *c*_1_ (i.e., for *t* ∈ (0, 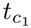 ]), we assume that the latent state evolves according to a *stochastic differential equation* (SDE) described by the Ornstein-Uhlenbeck (OU) process [55]. That is, if we let 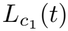 for *t* ∈ (0, 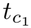 ] describe this latent state, then the OU model assumes that

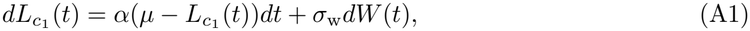

where *µ* ∈ ℝ is the latent state “optimum”, *α >* 0 determines the rate of reversion toward this optimum, and *σ*_w_ *>* 0 is the diffusion coefficient governing the magnitude of the stochastic fluctuations from *W* (*t*), a Wiener process. We similarly assume that 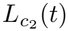 evolves independently from 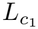 for 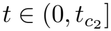 according to the same stochastic process. Then, we set

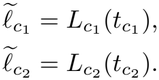

At times 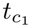 and 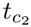, cells *c*_1_ and *c*_2_ respectively split into two cells each, and the latent states of their children will also follow the OU process described in Eq. A1. This process continues until time *t* = *t*_obs_, at which point the cells are observed.

While the SDE formulation of the OU process may feel a bit contrived in the context of cellstate evolution, by looking at the marginal distribution over ***l***, we can understand the OU process as encoding a kernel function of a Gaussian process (GP) that assumes an exponential decay (as a function of relatedness) in the covariation of the latent states across cells. That is, the OU process implies

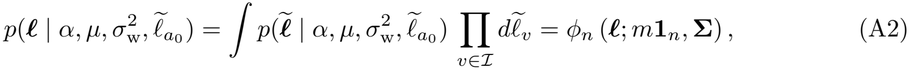

where *ϕ*_*n*_ represents the *n*-dimensional multivariate normal density,

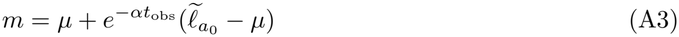

and

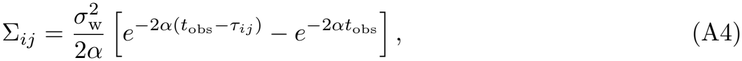

where *τ*_*ij*_ denotes the time from the root to the most recent common ancestor of leaves *i* and *j*.

When fitting the SOUR model in practice, care must be taken when setting the root state 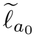 . Theoretically, 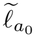 represents the gene expression state of a very distant ancestor, which is typically not possible to know with any degree of certainty. This might seem to suggest that instead we should place a prior over the root state and infer it from the data. However, due to the inclusion of an intercept term in our model (i.e., *β* in Eq. 5), this can lead to identifiability problems. To see this, it is easier to consider a reduced and slightly modified version of the SOUR model. Specifically, examine the model

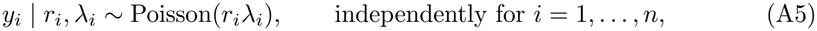

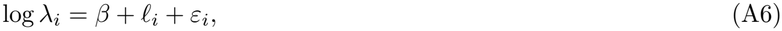

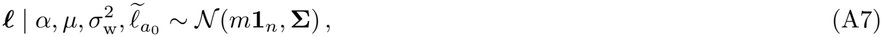

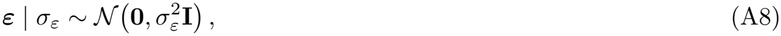

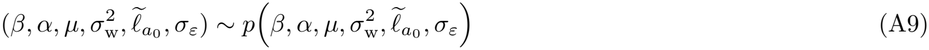

where *m* and **Σ** are as defined in Eqs. A3 and A4, respectively. The central simplification of this model relative to the SOUR model is that we have omitted the spatial term, while we have also allowed for a random root state 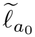 which induces a non-zero mean *m* on each element of ***l***.

To understand the difficulty with identifying *β*, 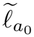, and *µ*, note that we can write the distribution of the log rates as

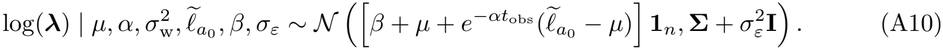

Then, for any constant *q* ∈ ℝ, adding *q* to both 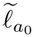 and *µ* and subtracting it from *β* will lead to

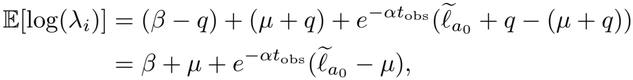

with covariances unchanged. To avoid the identifiability issues between *µ, β*, and 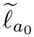, we must remove or constrain some of these parameters. In principle, if we removed *β* from the model, then both the root state 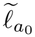 and *µ* are identifiable. However, because asymptotically the OU process has no “memory” of its root state (i.e., *β* + *m → β* + *µ* as *t*_obs_ *→ ∞*), this root state will generally be very weakly identified. Instead, we opt to consider the set of OU processes where 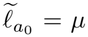 . In this case, the mean of the process is *µ* + *β* for any *t*_obs_. This further introduces an identifiability issue between *β* and *µ*, and to resolve this we can set *µ* = 0, which thus implies 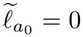 . Thus, by setting 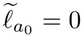 and just estimating *β*, we can represent any OU process that is started at its optimum.

In summary, to resolve the identifiability issue between the root, the intercept, and the OU optimum, we set *m* = 0 in Eq. A3 and use the covariance (modulo the additional identifiability constraint imposed by Eq. 13) in Eq. A4, with the re-parameterization

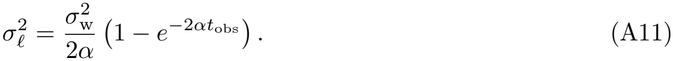

with this parameterization, the leaf covariance **Σ** of Eq. A4 is exactly **K**_*l*_ of Eq. 12.

#### A.3 Hyper-parameter Priors

The final piece required for our model is a prior distribution over the intercept term *β* and hyperparameters, ***ψ****s*, ***ψ****l*, and *σ*_*ε*_. We will use a prior distribution that factorizes over each of these terms

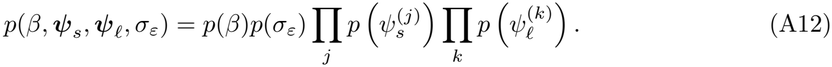

In the parameterizations discussed in Section 4.2, 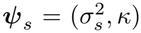, where 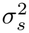 is the marginal variance of the exponential kernel and *κ* is its inverse length-scale. The precise form of ***ψ***_*l*_ depends on the experiment: in the full tree case, 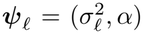, where 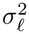 is the variance of the OU process at the observed cells (Eq. A11) and *α* is its reversion rate, and in the clonal case we simply omit the reversion rate, leaving 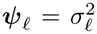, the variance of the clonal random effect. We write *σ*_*s*_ and *σ*_*l*_ for the corresponding marginal standard deviations, which is the scale on which the priors below are specified.

For our intercept term, we use a flat, uninformative prior:

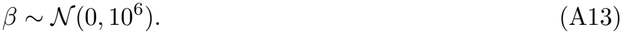

We also use a relatively uninformative prior for the variance 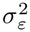,

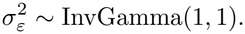

The remaining hyper-parameters control the two structured components. For many of their parameters, we use a *penalized-complexity* (PC) prior [56]. A PC prior is built around a *base model M*_0_: a simpler model, nested inside a flexible one, which is recovered by driving the hyper-parameter of interest to a limiting value. Writing *M* (*θ*) for the model in which that hyper-parameter takes the value *θ*, the extra complexity carried by *M* (*θ*) is measured by the Kullback–Leibler (KL) divergence of *M* (*θ*) from *M*_0_ [57],

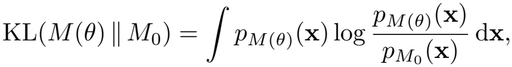

which is placed on the scale of a distance by taking

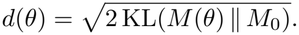

The prior is then specified on this distance as *d* ∼ Exponential(*ξ*). The exponential is used both because its mode is 0 (corresponding to a mode of *M*_0_), and because it corresponds to a constant rate of penalization in the space of the distance (i.e., *p*(*d* + *δ*)*/p*(*d*) = *r*^*δ*^ for some constant 0 *< r <* 1 determined by the setting of the exponential rate *ξ*).

For the marginal standard deviation parameters of both the spatial (*σ*_*s*_) and lineage (*σ*_*l*_) fields (generically *θ* = *σ*), the base model is the one in which the component standard deviation is set to 0 (*M*_0_ = *M* (0)). For a Gaussian component the distance *d*(*σ*) is proportional to the standard deviation *σ* [56], so the PC prior is an exponential on the standard deviation itself:

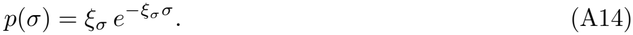

The rate *ξ*_*σ*_ is typically determined by making a statement of the form ℙ(*σ > U*_*σ*_) = *q*_*σ*_, and consequently setting 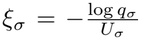 . The pair (*U*_*σ*_, *q*_*σ*_) can be thought of as specifying a soft upper bound *U*_*σ*_ on the standard deviation together with a tail probability *q*_*σ*_. Here, we fix *q*_*σ*_ = 0.05, and use a quasi-empirical-Bayes approach for determining *U*_*σ*_. Specifically, we fit a Poisson log-normal model that keeps only the fixed effects (i.e., Eqs. 4–9 with ***l*** = **s** = **0**; the intercept *β*, or the per-sample intercepts *β*_smp(*i*)_ of Appendix A.6, together with any covariates), obtain an estimate 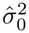 of the gene’s excess-Poisson variance, and set 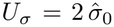 . By the properties of the exponential distribution, this implies

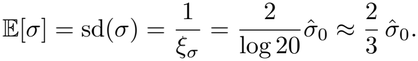

The same value of *U*_*σ*_ is shared by both the lineage and spatial marginal standard deviation priors, which act on *σ*_*l*_ and *σ*_*s*_ respectively.

The final two hyper-parameters that require priors are the length-scale parameters of both the OU component and the spatial component.

For the spatial field, it is conventional to work not with the inverse length-scale *κ* of Section 4.2.1, whose units are inverse distance, but with the spatial *range*

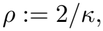

*≈*the distance at which the spatial correlation has decayed to *e*^*−*2^ 0.14. Previous work has already derived a PC prior on *ρ* [58], where the base model corresponds to *ρ* = *∞*, at which point the field becomes constant across the spatial domain. Using this base model, the distance *d*(*ρ*) is proportional to a negative power of the range, which in two-dimensional space is *d*(*ρ*) ∝ *ρ*^*−*1^. An exponential prior on the distance therefore makes 1*/ρ* exponentially distributed, and a change of variables gives

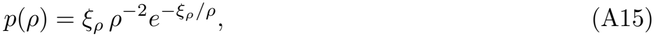

where *ξ*_*ρ*_ is the selected parameter of the exponential distribution on the distance. As with the standard deviation, the rate is fixed by a probability statement, though now one concerning the *lower* tail of the range, ℙ(*ρ < U*_*ρ*_) = *q*_*ρ*_, which gives *ξ*_*ρ*_ = *−* log(*q*_*ρ*_) *U*_*ρ*_. Because 1*/ρ* is exponentially distributed, *ρ* itself has a heavy right tail with no finite mean, and so the prior is most naturally summarized through its quantiles [58]. We set a weak prior of ℙ(*ρ <* 0.25) = 0.5, corresponding to a median range of 0.25 in the normalized spatial coordinates used in our experiments (see Appendix A.4.2 for more details on our standard normalization).

Despite the parallel between the reversion rate of the OU model (*α*) and the inverse length-scale of the spatial exponential prior (*κ*), the lineage length-scale admits no natural base model, and so no natural PC prior. For a fixed *t*_obs_, its two limits are *α t*_obs_ *→ ∞*, where 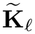 tends to 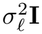 and the process becomes iid, and *α t*_obs_ *→* 0, where 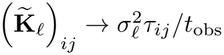, and the process becomes Brownian motion. Neither limit switches the lineage component off, nor does either limit produce a constant field, in the way that *ρ→∞* does for the spatial component, and so there is no direction along this axis for a PC prior to shrink toward.

Instead of a PC prior, we use an exponential prior directly on the dimensionless mean-reversion rate *αt*_obs_, taking

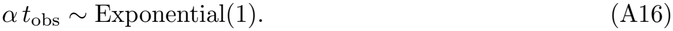

This prior discourages very large values of *α*, which correspond to OU processes that are very close to iid noise and thus difficult to separate from ***ε***. This prior also has the convenient interpretation under the OU model (Eq. A1) that the median time for a displacement from the optimum to decay to half its level is the height of the tree.

### A.4 Posterior Computation

Here, we describe our approach to approximating the posterior in Eq. 14 with the integrated nested Laplace approximation (INLA) [53]. To simplify notation, we will sometimes represent the collection of Gaussian latent variables as

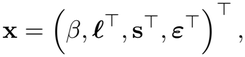

and represent the remaining covariance and variance parameters with the hyper-parameter vector

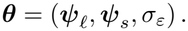

Using this notation, the posterior distribution of interest is

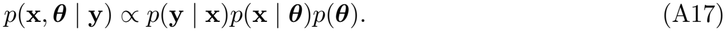

#### A.4.1 INLA Preliminaries

INLA is a method for approximating posterior distributions in the form of Eq. A17 when *p*(**x** |***θ***) is a multivariate normal distribution, which we write

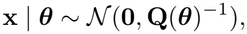

where **Q**(***θ***) is the precision matrix of **x**, which encodes conditional dependencies among its elements. The basic idea of INLA is to first approximate the posterior over the hyper-parameters as

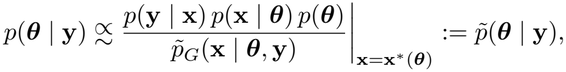

where 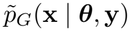 is a Laplace (Gaussian) approximation to *p*(**x** | ***θ*, y**), and

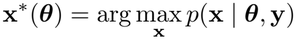

is the conditional posterior mode of the latent field for fixed ***θ***.

After fitting the model, INLA can generate approximate joint posterior samples from

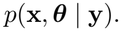

These samples are generated from a finite-mixture approximation.

##### Representing the SOUR Model as a GMRF

Because in the process of forming the posterior approximation INLA must repeatedly take Cholesky factorizations of **Q**(***θ***), the structure of this matrix can substantially affect runtime. In particular, if **x** has few conditional dependencies (i.e., **Q**(***θ***) is sparse), then these Cholesky factorizations are much less expensive. When **Q**(***θ***) is sparse, the model of Eq. A17 is said to be a *Gaussian Markov random field* (GMRF) [59].

The prior precision of the SOUR model has the form

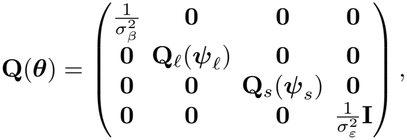

where 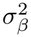 is the prior variance of *β* (which we fix as described in Appendix A.3), and **Q**_*ℓ*_(***ψ***_*ℓ*_) ≔ **K**_*ℓ*_(*T* ; ***ψ***_*ℓ*_)^*−*1^ and **Q**_*s*_(***ψ***_*s*_) := **K**_*s*_(**C**; ***ψ***_*s*_)^*−*1^ are the prior precision matrices over the lineage and spatial random effects, respectively. Here, both **Q**_*ℓ*_(***ψ***_*ℓ*_) and **Q**_*s*_(***ψ***_*s*_) are dense, *n × n* matrices, which greatly slows down computation in INLA.

To handle the dense spatial precision matrix **Q**_*s*_(***ψ***_*s*_), we exploit the fact that the exponential kernel of Section 4.2.1 is exactly the Matérn covariance with smoothness *ν* = 1*/*2, and therefore admits the widely used SPDE representation of [60]. Specifically, the field is approximated by a basis expansion over a triangulated mesh of the tissue, whose coefficients form a GMRF with sparse precision, replacing direct computation with the dense **K**_*s*_ by local interactions between neighboring mesh vertices. This construction is exact only when *ν* +*d/*2 is an integer, which for our field (*ν* = 1*/*2 in *d* = 2 dimensions) equals 3*/*2; for such non-integer values we use the approximation supplied by INLA [53, 60].

In practice, several parameters must be set to construct a mesh for a given dataset. For each of our experiments, we first normalize the spatial coordinates into [0, 1] *×* [0, 1] while preserving the aspect ratio of the data (for the SpaceBar dataset we do this separately for each sample, and for the PEtracer dataset we do this for the combined coordinates of all three samples). In all of our experiments, we build our mesh by first constructing a non-convex hull around the spatial domain (by calling `fm_nonconvex_hull` from the `fmesher` package [61] with `convex` = -0.03, `concave` = 0.03 and `resolution` = 100). See the orange line of Fig. A2 for an example.

We then build a mesh inside (and an extension around) this hull by calling `fm_mesh_2d_inla`. The extension is needed because the SPDE representation inflates the variance of the field near the edge of the mesh, and thus expanding the mesh beyond the data prevents this effect from influencing the estimated spatial field at the observed points [60, 62]. Building the mesh then requires setting the `max.edge` parameter, which accepts a pair of values that set the longest edge that any triangle of the mesh is permitted to have (the first argument for the mesh within the hull, the second for the mesh outside). The size of the triangles determines the length-scales resolvable when fitting the model: variation at scales much smaller than the triangles cannot be captured. However, computational complexity increases with the size of the mesh. In all our real-data analyses and simulations, we set `max.edge` to a fixed physical length. Specifically, writing *q* for the number of microns spanned by one unit of the normalized coordinates of a given sample, we take

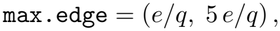

so that an interior triangle edge always spans the same physical distance *e*, whatever the extent of the sample. We use *e* = 75 *µ*m, or approximately 6.6 cell diameters, for both the SpaceBar and PEtracer datasets, which we expect to be a reasonable default for other datasets in practice. We show an example mesh in Fig. A2.

To handle the dense lineage precision matrix **Q**_*ℓ*_(***ψ***_*ℓ*_), we employ an expanded (but exact) parameterization of the lineage field. Specifically, while the prior precision matrix over ***ℓ*** is dense, the prior precision matrix over 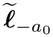 (i.e., all vertices of the tree except the fixed root 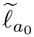) is extremely sparse.

**Fig. A2.**
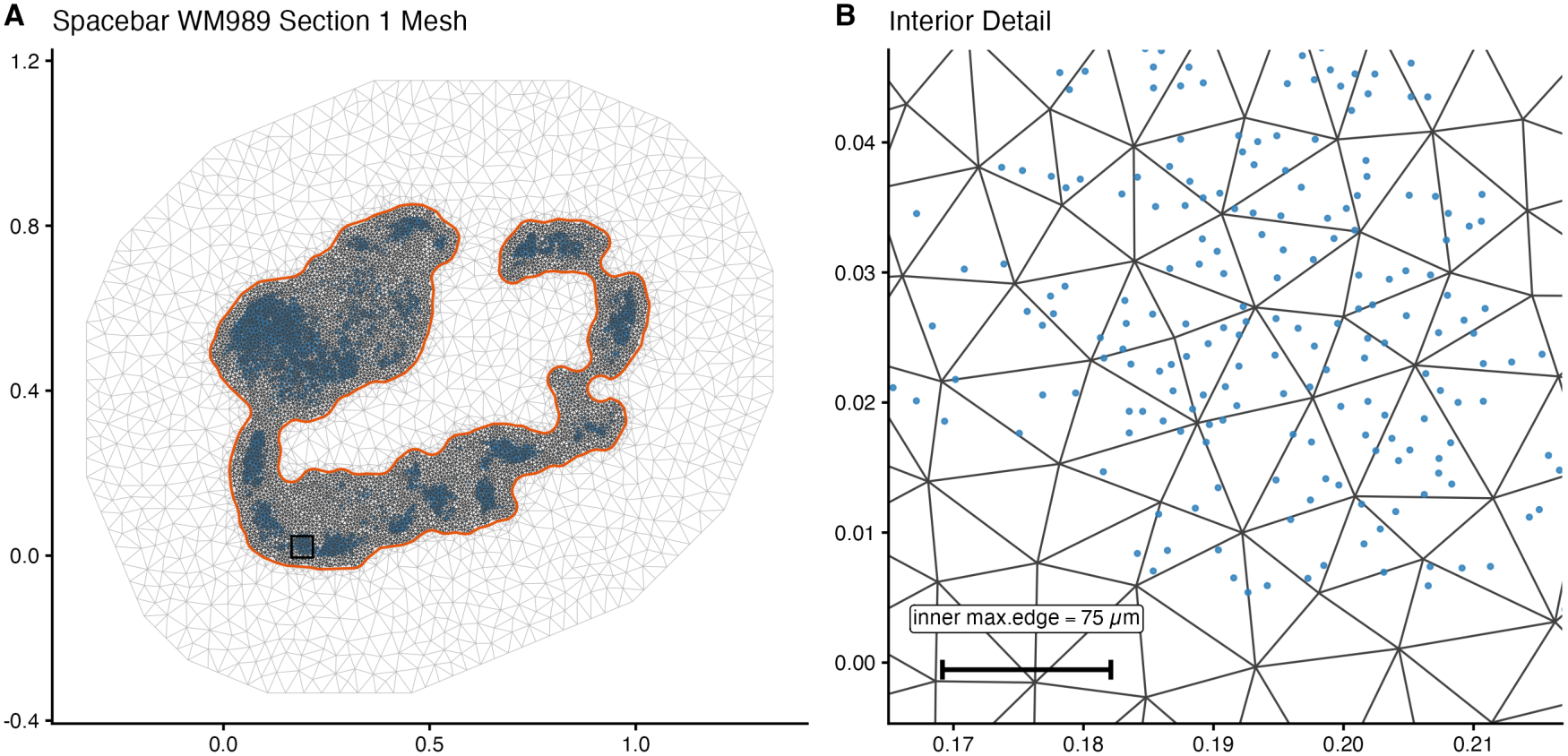
The mesh used for the SPDE representation of the spatial field, shown on sample 1 of the SpaceBar dataset. **(A)** The whole sample. Cells are blue, the non-convex hull is orange, mesh triangles inside the hull are dark and those of the exterior extension are light. The box marks the region enlarged in **(B)**, a patch that lies in the interior.

To see this, recall that by construction any two sibling nodes 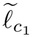 and 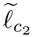 are conditionally independent given the state of their parent 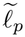. Thus, the only non-zero off-diagonal entries of **Q**_*ℓ*_(***ψ***_*ℓ*_) correspond to parents and children. While this introduces more parameters for INLA to estimate a posterior over, it replaces a dense *n × n* precision matrix 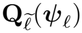 with a sparse (|*V*| *−* 1) *×* (|*V*| *−* 1) matrix 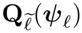 (where *V* and *ℰ* are the vertex and edge sets of the tree as above) carrying on the order of |*ℰ*| *≪ n*^2^ non-zero entries.

#### A.4.3 Poisson Log-Normal Likelihood

Relative to asymptotically exact MCMC, we found that the posterior approximation to the iid latent effect ***ε*** (and to its variance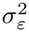) performed poorly. To mitigate this issue, we decided to instead fit INLA to the model with ***ε*** integrated out. This is equivalent to replacing the Poisson likelihood on each *y*_*i*_ with a Poisson log-normal likelihood

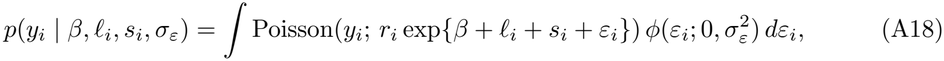

where *ϕ* denotes the normal density. Writing *ζ*_*i*_ = *β* + *ℓ*_*i*_ + *s*_*i*_ for the *structured* linear predictor, we denote this marginal likelihood by 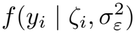. The integral of Eq. A18 has no closed form, but it can be evaluated efficiently and accurately using adaptive Gauss–Hermite quadrature (AGHQ) [63]. Fitting INLA to this marginal likelihood means that the iid effect ***ε*** is no longer part of the latent field, and thus in the fitting step we estimate the posterior only over **x**_*−****ε***_ =( *β*, ***ℓ***^*⊤*^, **s**^*⊤*^) ^*⊤*^ together with the hyper-parameters ***θ***.

Because ***ε*** is integrated out of the likelihood, a given posterior sample *j* returned by INLA contains 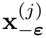 and the hyper-parameters ***θ***^(*j*)^, but not the iid effects. This prevents us from computing posterior quantiles of the variance shares (e.g., Eq. 15), which are functionals of the *complete* field realization, including ***ε***. We therefore augment each INLA sample by drawing each *ε*_*i*_ from its one-dimensional full conditional given the sampled structured predictor 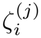 and iid variance 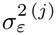,

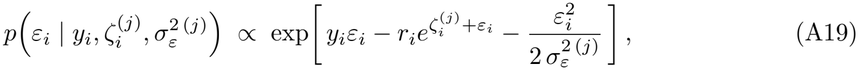

whose normalizing constant is precisely the marginal likelihood in Eq. A18 evaluated at the sampled values. We draw each 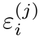 from this conditional exactly by rejection sampling.

### A.5 Additional Details on Generation of Simulated Data

#### A.5.1 PEtracer Simulations

In the spatial-only PEtracer simulations, we generated 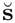 from the exponential covariance assumed by our model (Eq. 11), with *κ* = 0.003817 *µ*m^*−*1^. We selected this value of *κ* because it made the half-life of the spatial signal, defined as the distance at which the correlation between two cells falls to 1*/*2, equal to approximately 16 cell diameters (assuming each cell is 11.35 *µ*m in diameter, which is the average diameter across the PEtracer and SpaceBar datasets). This value of *κ* falls squarely within the distribution of fitted values for the PEtracer dataset (Fig. B5A). We show simulation results with other values of *κ* in Fig. B4C.

In the lineage-only PEtracer simulation, we generated 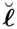 from a Brownian Motion (BM) model, instead of the OU model assumed by SOUR. This model implies 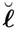 is drawn from a mean **0** multivariate normal with a covariance dictated by the BM process. The unconstrained covariance can be derived by taking the limit *α →* 0 of Eq. 12 at a fixed tip variance 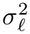 (Eq. A11), yielding

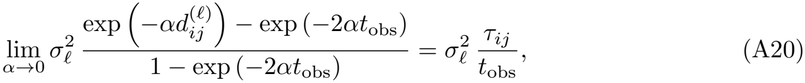

where the last equality uses 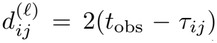. We then apply the formula Eq. 13 to Eq. A20 before generating data. Because our prior for *α* (see Appendix A.3 for more details) only has mass on *α >* 0, this BM covariance cannot be represented exactly by the SOUR model. This covariance implies a lineage half-life of 35 days, which is the height of the phylogenetic tree.

#### A.5.2 SpaceBar Simulations

For the SpaceBar spatial-only simulations, we again set *κ* = 0.003817 *µ*m^*−*1^. However, instead of defining 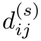 as the Euclidean distance, as the SOUR model assumes, we set

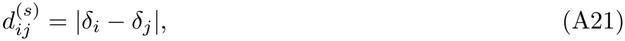

where *δ*_*i*_ is the Euclidean distance of cell *i* from the boundary of the tumor (i.e., the shortest distance from **c**_*i·*_ to any point outside the tumor). This construction causes a radial gradient in gene expression, and was inspired by an observation in the SpaceBar paper that the genes with the greatest spatial variability tended to show a gradient of expression between the interior and exterior of the tumor. Fig. C15 shows the tumor boundary for the SpaceBar sample we use for the simulations, as well as an example spatial field when using this distance metric. We show the distribution of fitted values for *κ* in the SOUR model (with kernel defined by the Euclidean distance) in Fig. B5C, and show additional simulation settings of *κ* in the radial kernel in Fig. B4A.

Finally, for the lineage-only simulations, we generate data exactly as described in Section 4.2.3.

### A.6 Modification of SOUR Model for PEtracer Real-Data Analysis

The PEtracer dataset analyzed in Section 2.5.2 contains three samples from a single mouse. The cells from these samples all lie on a single lineage tree, but the spatial coordinates across samples are not naturally on the same coordinate plane. In principle, as a pre-processing step we could attempt to align the samples onto a shared coordinate plane using, for example, PASTE [64]. However, the shape of the tumor appears to change to some degree across the samples (see Fig. 5E–G), and thus there may be ambiguity in exactly how to align the samples. Moreover, we do not have exact information about the physical distance in the original tumor between each sample, and thus it is unclear how to calculate the distance between cells across samples, even if their x-y coordinates were aligned.

We therefore fit a variant of the SOUR model in which each sample has its own intercept and spatial field, while the lineage field is shared across samples through the common tree. Writing smp(*i*) *∈ {*1, 2, 3*}* for the sample containing cell *i, n*_*k*_ for the number of cells in sample *k*, and idx(*i*) *∈ {*1, …, *n*_smp(*i*)_*}* for the position of cell *i* within its own sample, we assume

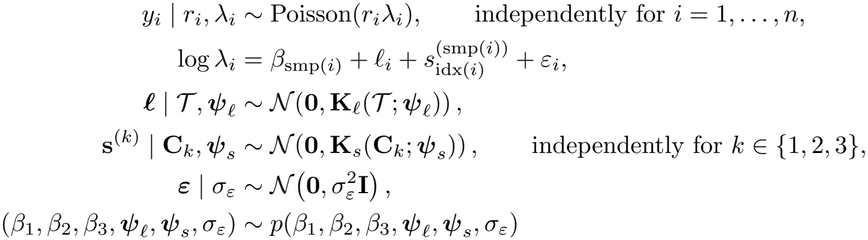

The three spatial fields are thus independent *a priori* but share the hyper-parameters ***ψ***_*s*_, so every sample informs a common spatial variance and length-scale. All hyper-parameters have the same priors as described in Appendix A.3, and we assume each *β*_*k*_ is drawn independently from a *N*(0, 10^6^), as in Eq. A13. As in the single-sample model, each spatial field **s**^(*k*)^ is constrained to have mean zero over the cells of its own sample, by applying Eq. 11 to **K**_*s*_(**C**_*k*_; ***ψ***_*s*_) separately for each *k*; no field is constrained at the cell positions of another sample.

### A.7 Selection of Genes for 4T1 Analyses

A common problem with imaging-based spatial transcriptomic data is that, due to the necessary pre-processing step of cell segmentation, some transcripts are misassigned to neighboring cells [65]. While this likely occurred in both the SpaceBar and the PEtracer datasets, the segmentation errors will have a different effect in the two analyses due to differences in experimental design. Specifically, the SpaceBar dataset is a chimeric system, where the tumor cells originate from a human but the stromal cells originate from a mouse. Because the SpaceBar gene panel was designed only to detect human genes, the only spillover that will occur is between tumor cells. The likely consequence on the SOUR model, relative to if the data had been segmented perfectly, is that the noise share will increase slightly and the structured shares will decrease slightly. In contrast, the PEtracer dataset is derived from an autochthonous mouse model, where the tumor and stromal cells are from the same organism. The PEtracer panel contained genes highly expressed in tumor cells, as well as other genes more highly expressed in stromal cells. As a result, spillover in the PEtracer dataset can occur between both tumor cells and surrounding healthy tissue. In the SOUR model, this could introduce spuriously large spatial shares in cases where the normal cell type that most highly expresses a gene is spatially organized. Fig. A3 provides an example of this effect, where *Arg1*, a gene expressed at only very low levels in malignant cells based on the scRNA-seq data, is highly expressed in tumor cells that are close to the cell type annotated “Arg1 macrophages”.

The MERFISH panel used for the PEtracer dataset [21] measured 175 genes in the three samples analyzed here. In addition to the spatial lineage tracing data, [21] released cell-type-annotated dissociated single-cell RNA-sequencing (scRNA-seq) data of 4T1 tumor cells and surrounding stroma. We used this scRNA-seq data as a reference to determine genes that had suspiciously high expression in the malignant cells in the MERFISH samples. All reference comparisons use the log-normalized expression values released with the data (the natural logarithm of one plus the library-size-normalized count). Specifically, we retained genes in the panel if at least one of the following held:

1. The mean log-normalized expression across malignant cells of the tumor sample was at least 0.1.
2. The mean log-normalized expression in malignant cells was higher than the mean in every other annotated cell type (the immune, endothelial and fibroblast populations), pooling cells from the tumor and lung samples.

We additionally excluded any genes that were not expressed in at least 5% of cells in each sample, and the gene *Arg1*, whose mean log-normalized expression in the reference was 22-fold higher in macrophages than in malignant cells of the tumor sample (roughly 100-fold on the expression scale).

This left 39 genes (see Fig. C13 for the complete listing).

### A.8 Marginal Range Method

Here, we define mathematically the marginal range method originally introduced in the SpaceBar publication [23]. Let *P* = *{ A*_1_, …, *A*_*K*_*}* denote a partition of the *n* cells into *K* non-overlapping groups. For the clone range, these partitions are defined by clonal identities, and for the spatial range, these partitions are defined by spatial regions determined by the original authors. For a given partition, define the mean expression of the given gene in group *A*_*k*_ as

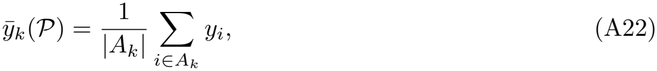

and define the corresponding partition-level expression range as

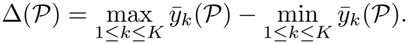

**Fig. A3.**
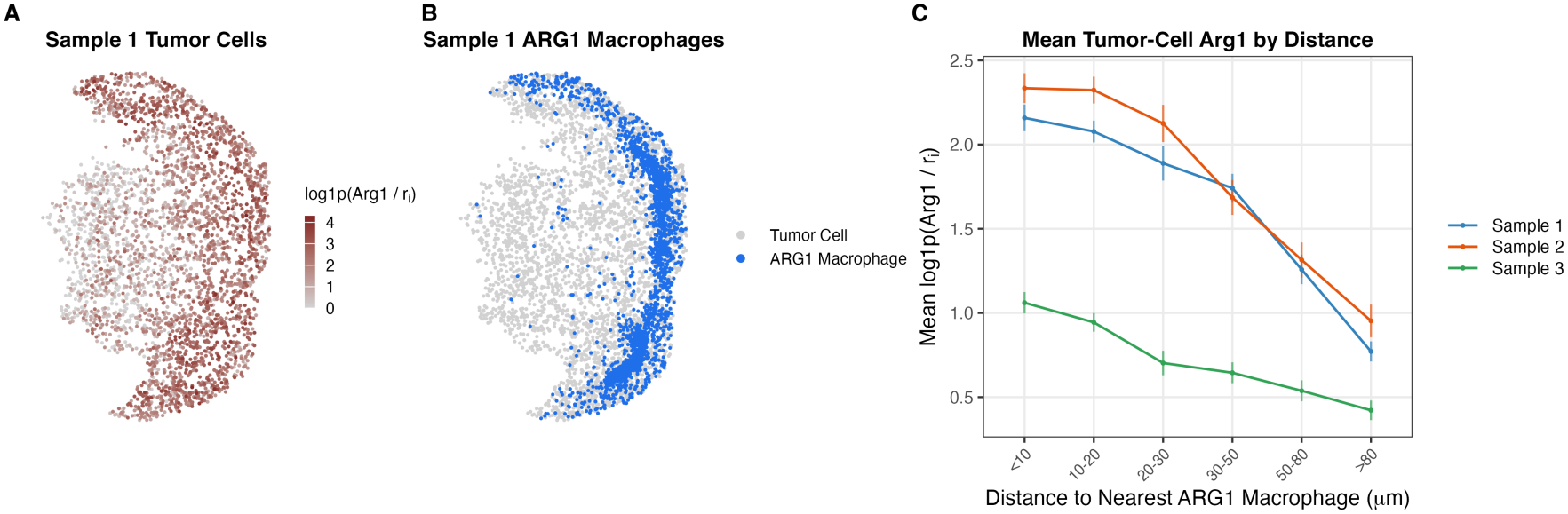
Example of segmentation errors in the PEtracer dataset, where *r*_*i*_ is the scaling factor for cell *i* defined in Section 4.2. **(A)** Normalized expression of *Arg1* in tumor cells of sample 1 of the dataset. **(B)** Sample 1 tumor cells and Arg1 macrophages, as annotated in [21], colored by cell type. **(C)** Average normalized expression of *Arg1* in tumor cells as a function of distance to nearest Arg1 macrophage. Vertical lines are 95% confidence intervals for the bin mean.

To correct for the fact that a non-zero range is expected even under no association between expression and the partition, the observed range is normalized by a permutation-based null range. Let *P*^(*m*)^, for *m* = 1, …, *M*, denote the partition obtained by randomly permuting the group labels across cells while preserving the observed group sizes. Define

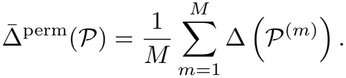

The normalized partition score is then

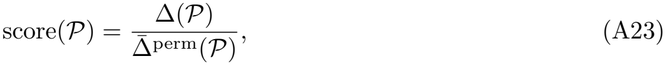

so that a value of 1 indicates a range equal to what is expected under the null. *p*-values are then computed by comparing the observed range to its permutation distribution:

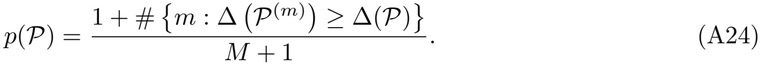

In the original SpaceBar publication [23], Eq. A22 is computed on the raw gene expression values, as written. We use the published scores and *p*-values for each gene whenever making comparisons between SOUR and the marginal range method on the SpaceBar dataset. However, when applying this method to the raw counts in our simulated SpaceBar datasets, we found that even when all gene expression variation was caused by noise (i.e., *v*_*ε*_ = 1), the *p*-values of this method were not properly calibrated. Further investigation revealed that this was because the size factors, *r*_*i*_ for *i* = 1, …, *n*, had spatial structure, in that some regions of the tissue had more total gene expression counts per cell than others. Because Eq. A22 does not account for the size factors *r*_*i*_, this caused anti-conservative *p*-values for both the spatial and clonal scores. We felt that this miscalibration gave an unfair advantage to SOUR, which includes the values of *r*_*i*_ used to generate the data explicitly in the model (Eq. 4).

To make our simulated comparisons more fair, we instead computed Eq. A22 using Pearson residuals from a correctly specified null model. Specifically, under the hypothesis that a gene carries neither spatial nor lineage structure, the model of Eqs. 4–8 reduces to

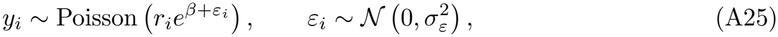

independently across cells. Writing 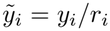, under Eq. A25 we have 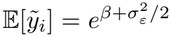, and setting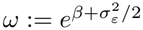, we can write

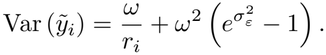

Letting 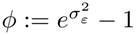, we can derive the following moment based estimators:

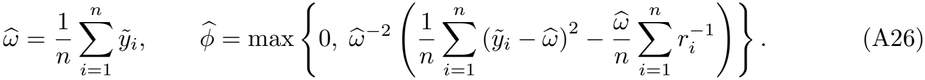

The Pearson residual of cell *i* is then

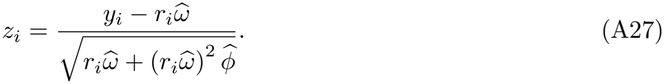

By construction *z*_*i*_ has mean 0 and variance 1 for every cell *i*. In our simulations, we therefore replaced *y*_*i*_ with *z*_*i*_ in Eq. A22 and left every other aspect of the method unchanged.

### A.9 Marginal Autocorrelation Method

The marginal autocorrelation method used in the original PEtracer publication [21] is based on a Moran’s I statistic [66]. Specifically, given observations *x*_1_, …, *x*_*n*_ with mean 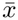 and a matrix of non-negative weights **W** encoding how close each pair of observations is (with *w*_*ii*_ = 0), Moran’s I is defined as

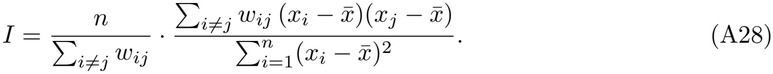

The PEtracer publication uses binary weight matrices, **W**^(*l*)^ and **W**^(*s*)^ for the marginal lineage autocorrelation and the marginal spatial autocorrelation, respectively. For the lineage component, the original authors set 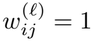 if cells *i* and *j* shared a common ancestor in the last 10 days (based on the reconstructed lineage tree) and 0 otherwise. For the spatial component, 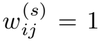 if cells *i* and *j* are within 100*µ*m, and 0 otherwise. The autocorrelations are then computed for lineage and space on the raw counts **y** for each gene (the original authors note that they compute Eq. A28 on the z-scored expression values, but Moran’s I is invariant to shifting and scaling a vector).

For all real-data results on the PEtracer dataset, we display the published autocorrelation values, computed as described above. However, for the PEtracer simulations, we make two modifications to this procedure. First, we compute autocorrelations on the vectors of Pearson residuals **z** (Eq. A27), computed for each gene as described in Appendix A.8. Second, instead of defining nearest neighbors for the weight matrices **W**^(*l*)^ and **W**^(*s*)^ based on the criteria described above, we define these matrices so that *w*_*ij*_ = 1 if cell *j* is in the 10 nearest neighbors of cell *i* (where nearness is defined 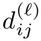 for 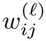 and by 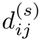 for 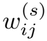. We make this choice because our simulations only use one of three samples from the PEtracer dataset, and thus the number of cells with a common ancestor within the last 10 days is greatly reduced relative to the original dataset. Using 10 nearest neighbors increases the number of neighbors, and for the sake of symmetry we use the same construction for the spatial weights. We do not expect that our results on the confounding of the marginal autocorrelation method are directionally sensitive to the choice of nearest neighbor construction.

### A.10 Runtime and Computing Environment

#### Hardware and software environment

All model fits reported in this paper were run on a single workstation containing one Intel Xeon W-2295 processor. The processor has 18 physical cores and 36 hardware threads. The machine ran Ubuntu 22.04.5 (Linux kernel 6.8.0), R 4.5.1 and R-INLA 25.10.19.

#### Runtime

Because the SOUR model is fit separately for each gene, the computation parallelizes trivially across genes. We therefore ran each fit single-threaded and fit 36 genes concurrently, one per hardware thread.

On the SpaceBar data, fitting all 108 genes for sample 3 (15,226 cells, 59 clones, and a spatial mesh with 4,739 nodes) took 8 minutes and 44 seconds, with a median of 2.1 minutes per gene (range 1.2–3.9 minutes).

The PEtracer analysis is considerably more expensive, because its lineage field is defined on a tree with 16,889 nodes shared across all three samples. Fitting all 39 genes for the combined three-sample model (10,061 cells, 16,889 tree nodes, and 1,683 spatial mesh nodes) took 3 hours and 9 minutes, with a median of 2.1 hours per gene (range 0.9–3.1 hours).

## Appendix B Supplementary Results

### B.1 Simulation Results

#### B.1.1 Extended Main Text Simulations

As described in Section 4.5, the simulations in Fig. 3 are presented for only one choice of spatial half-life and lineage half-life. In Fig. B4, the results of Fig. 3 are shown alongside other choices of the half-life parameters that are representative of the fitted half-lives in our real-data analyses (Fig. B5).

Across values of both the lineage and spatial half-lives, SOUR assigns negligible variance shares to the non-driving signal, and thus remains robust to the correlation between spatial and lineage distances. However, for smaller half-life values, SOUR appears to give downwardly biased estimates of the variance shares for the driving signal, and correspondingly upwardly biased variance shares for the noise component. Because the covariance of the driving signal approaches that of iid noise as its half-life shrinks, becoming exactly iid in the limit, the conservative PC priors placed on the spatial and lineage variance terms (Appendix A.3) increasingly favor the iid component over the structured one. In principle, this bias could be attenuated by placing a less conservative prior on the structured variance parameters. However, in a scenario where all expression variation were driven by unstructured noise, using less conservative priors could lead to inflated variance shares for the structured components, as nothing in the likelihood encourages the model to favor the unstructured component. We view the conservative behavior of the SOUR model to be preferable to falsely attributing noisy variation to structured signals, which could lead to wasted effort in functional follow-up studies.

Finally, in Fig. B6, we provide results of fitting the SOUR model to data generated with the same covariance structures as described in Section 4.5, but with contributions from both space and lineage in each dataset. Specifically, we set 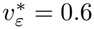, and varied 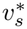 and 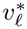 on a grid such that 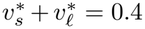. We performed these experiments on the same grid of half-lives for space and lineage as Fig. B4.

For signals with longer half-lives, SOUR reports extremely accurate variance shares (e.g., the PEtracer simulation with a 35-day lineage half-life and 64-cell spatial half-life; Fig. B6A). However, for signals with shorter half-lives there is again a downward bias of the structured shares toward the iid component.

#### B.1.2 Additional Simulations Testing Robustness of SOUR to Model Misspecification

We created two additional simulation studies to further test the robustness of SOUR’s variance shares to model misspecification. In these simulations, we still generated data from the additive structure assumed by SOUR, as described in Section 4.5. However, we varied the form of the covariance matrices used to generate the lineage and spatial fields, as described below.

##### Lévy simulation

The PEtracer lineage-only simulation (Fig. 3B) presents the results of SOUR fit to lineage data generated with a BM covariance, as described in Appendix A.5.1. While this in itself represents a form of model misspecification for SOUR, which assumes an OU model for the lineage component, the SOUR model can get arbitrarily close to a BM model by fitting an extremely small value of *α*. To increase the degree of model misspecification, here we generated data from a Lévy process. The Lévy process is a model of punctuated evolution, in which traits change in occasional bursts rather than gradually [67, 68]. Concretely, along a branch of length Δ*t*, the lineage process accumulates an increment

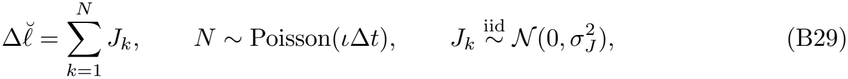

and the value 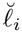 at a leaf is the sum of these increments along the path from the root to cell *i. ι* (the jump rate) and 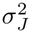 are hyper-parameters of the Lévy process, and under this process

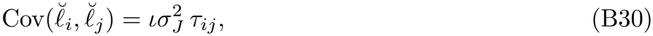

where *τ*_*ij*_ is defined as in Section 4.2.2. This covariance is of the same functional form as the BM covariance (Eq. A20); however, the distribution of 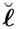 is not multivariate normal. Thus, generating data from a Lévy process tests the robustness of SOUR to non-normality of the lineage process.

**Fig. B4.**
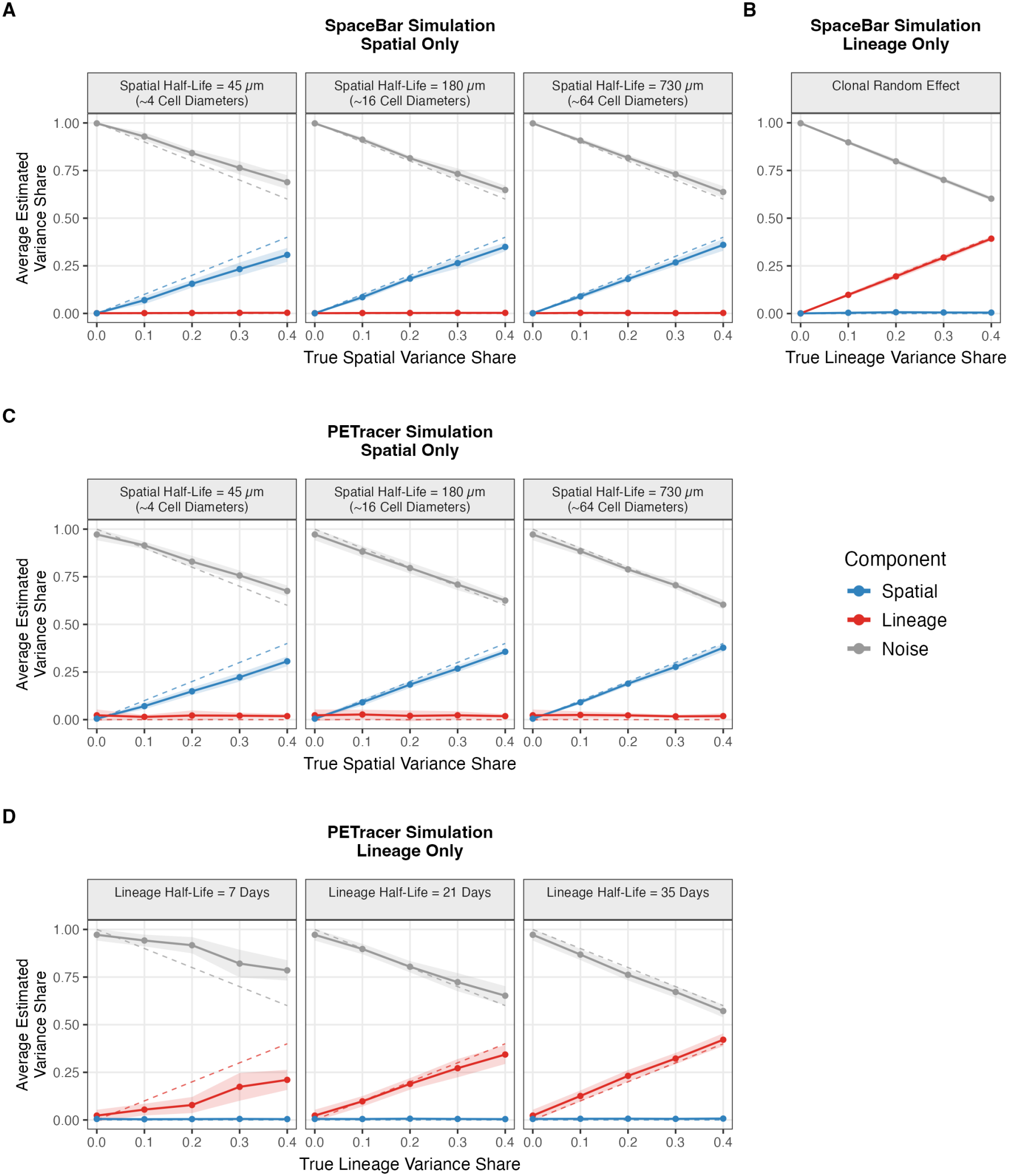
SOUR posterior mean variance shares on the same simulated data as Fig. 3, across more simulated length-scales. **(A)** SpaceBar spatial-only, **(B)** SpaceBar lineage-only, **(C)** PEtracer spatial-only, **(D)** PEtracer lineage-only. Fig. 3 reports the 16 cell half-life column of (A) and (C), the single panel of (B), and the longest-half-life column of (D), which is the Brownian motion limit of Eq. A20. Dashed lines indicate the true variance shares of the components. Each point is the mean over 25 replicate datasets, with bands indicating *±*1 SD across them.

In our simulations, we first generated 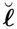 as described above, then we applied the linear map

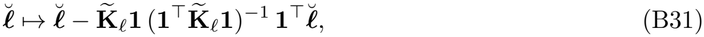

taking 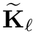 to be the BM covariance of Eq. A20, before normalizing 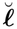 to its target variance share as described in Section 4.5. This map forces 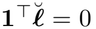, so that the simulated lineage field obeys the same identifiability constraint as the field that SOUR fits. Here, we set 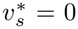 and used the same grid of values for 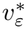 and 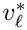 as Fig. 3, and generated data using sample 3 of the PEtracer dataset.

SOUR gave excellent performance in the Lévy process simulations. Even when the data departed greatly from normality (Fig. B7B), SOUR still estimated the variance shares quite accurately (Fig. B7C).

##### Paciorek-Schervish simulation

Finally, we test the robustness of the SOUR model to non-stationarity of the spatial process. To do this, we adapted the non-stationary Matérn kernel introduced by Paciorek and Schervish [70, 71], which we refer to as the *PS kernel* for short.

The PS kernel generalizes the Matérn kernel to have a length-scale that varies across space. Here, we set the Matérn smoothness parameter to 1*/*2 (matching the exponential kernel of Eq. 10), and define a varying length-scale as a function of the second coordinate of each cell. It is most convenient to write the resulting kernel in terms of a *local spatial range ρ*_*i*_ at each cell *i*, where *ρ* = 2*/κ* is the spatial range introduced in Appendix A.3. At smoothness 1*/*2, the PS kernel is

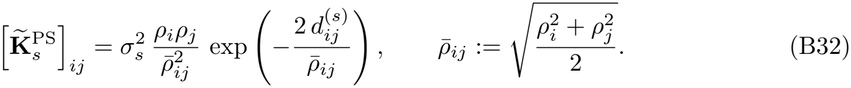

When *ρ*_*i*_ = *ρ* for every cell, Eq. B32 reduces exactly to the stationary exponential kernel of Eq. 10.

To create a gradient of length-scales across the tissue, we set the local range to vary log-linearly along the second coordinate axis:

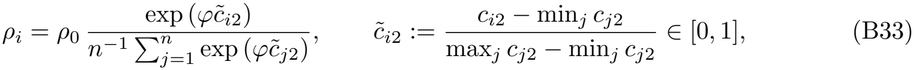

where *φ ≥* 0 sets the strength of the gradient and the denominator rescales the local ranges so that 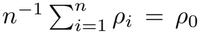 for every value of *φ* (see Fig. B8B to see how the half-life changes across the tissue for different values of *φ* and Fig. B8A for how this changes the generated spatial field). Note that *φ* = 0 recovers the stationary kernel. We set *ρ*_0_ = 0.2 in the normalized spatial coordinates, corresponding to a spatial half-life of approximately 14 cell diameters, and generated data with *φ ∈ {*0, 2.5, 5, 7.5*}* . As in our other simulations, we apply Eq. 11 to Eq. B32 before generating 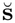, so that the simulated spatial field satisfies the same sum-to-zero constraint as the field that SOUR fits. We generated spatial-only simulations using this scheme on sample 3 of the PEtracer dataset, using the same settings of the value of 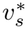 as in Fig. 3.

SOUR remains robust to the correlation between spatial and lineage distance across values of *φ* (Fig. B8C). However, SOUR displays a consistent downward bias of the spatial share and corresponding upward bias of the noise share for non-stationary kernels, and this bias appears to increase as the degree of non-stationarity increases (i.e., as *φ* increases; Fig. B8C). We again suspect that this may be a result of the PC priors on the parameters of the exponential kernel in the SOUR model (Appendix A.3). Specifically, as *φ* increases, much of the variation in the tissue looks like unstructured noise, and so perhaps the likelihood favors the noise component in these scenarios.

### B.2 Evaluation of Poisson Log-Normal Likelihood

In Appendix A.4.3 we noted that INLA approximates the posterior of the iid latent effect ***ε*** poorly, which motivated our decision to integrate ***ε*** out of the likelihood. Here, we demonstrate that our approach indeed yields more accurate posterior approximations (relative to MCMC).

While ideally we would compare our posterior approximation approach to MCMC on our real data analyses, estimating a posterior distribution using MCMC to these large datasets is quite computationally intensive. Instead, we compare the approximations on smaller, simulated datasets, and just focus on estimation of the lineage and noise components.

Specifically, in each of our simulations we generated data from the SOUR model on a perfect binary tree with 1024 leaves, setting 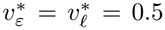. We set *t*_obs_ = 1 and *α* = 1, and set *r*_*i*_ = 1 for all cells. Because we suspected that the sparsity of the data would influence the quality of the INLA posterior approximations, we varied the value of the intercept, *β*, on a grid that resulted in some simulations with very sparse data and others with few zeros. To increase the speed with which models were fit, we omitted the spatial term **s** from the model.

To each of 30 replicate simulated datasets for each value of *β*, we fit the model described above using Hamiltonian Monte Carlo [72], INLA with the Poisson log-normal likelihood of Eq. A18, and INLA with an explicit approximation to ***ε***.

Overall, integrating out the iid effect improved the quality of the INLA approximation to the posterior of the lineage variance share. The explicit iid effect over-estimates the lineage variance share relative to MCMC in every one of the 180 replicates, by between 0.024 and 0.092 depending on the count level, whereas the error of the Poisson log-normal likelihood is smaller at every value of *β* and changes sign across the grid (Fig. B9A). Relative to the posterior standard deviation of the MCMC fit, the error of the explicit effect is roughly flat, between 0.32 and 0.45 SD, while the error of the Poisson log-normal likelihood ranges from 0.13 to 0.35 SD (Fig. B9B). Comparing the posteriors in full rather than only their means gives the same ordering: the 2-Wasserstein distance to the MCMC posterior is smaller for the Poisson log-normal likelihood in 161 of the 180 replicates, with a mean of 0.046 against 0.067 for the explicit effect (Fig. B9D).

The Poisson log-normal likelihood does, however, understate the uncertainty in the variance share. Both INLA fits report narrower credible intervals than MCMC at all but the sparsest count level, and the Poisson log-normal intervals are the narrowest everywhere (Fig. B9C). Pooled over all replicates, this costs some coverage of the true variance share, which is 88% for the Poisson log-normal likelihood, against 94% for MCMC and 95% for the explicit effect. We consider this an acceptable trade for the substantially more accurate point estimates, particularly because the bias of the explicit effect is in the same direction in every replicate.

**Fig. B5.**
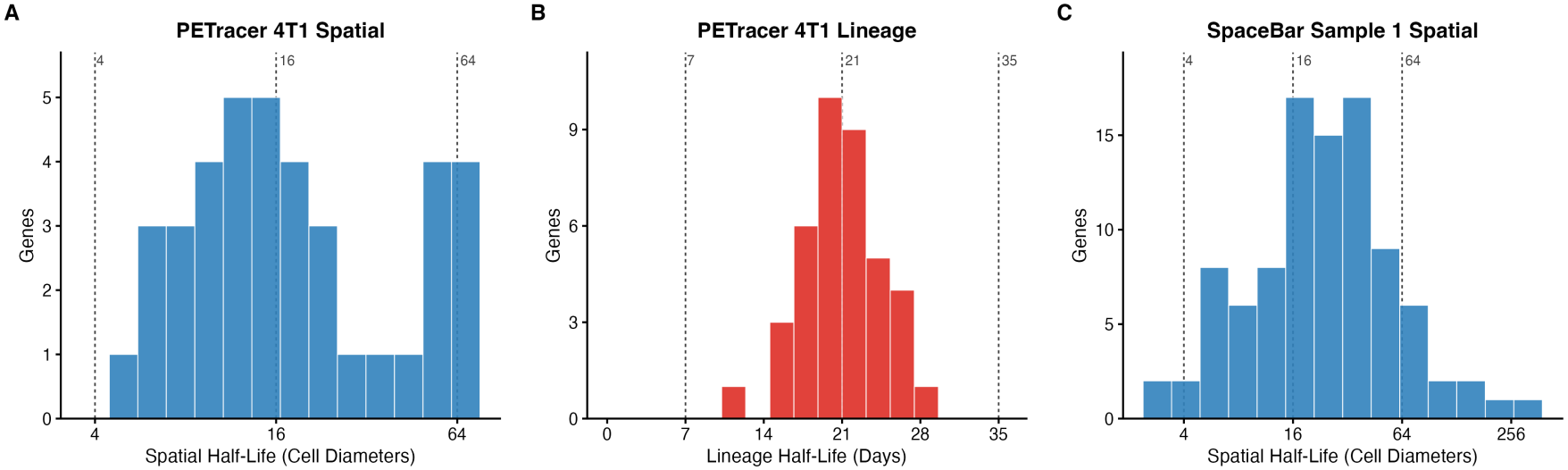
Histogram of posterior means of half-life parameters for the PEtracer and SpaceBar re-analyses. **(A)** Spatial half-life, log(2)*/κ*, of the PEtracer fit, in units of cell diameters. **(B)** Lineage half-life, 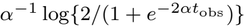, of the same 39 genes, in days. **(C)** Spatial half-life of the SpaceBar sample 1 fit (96 genes), drawn as in (A). Dashed vertical lines mark the simulated half-lives.

**Fig. B6.**
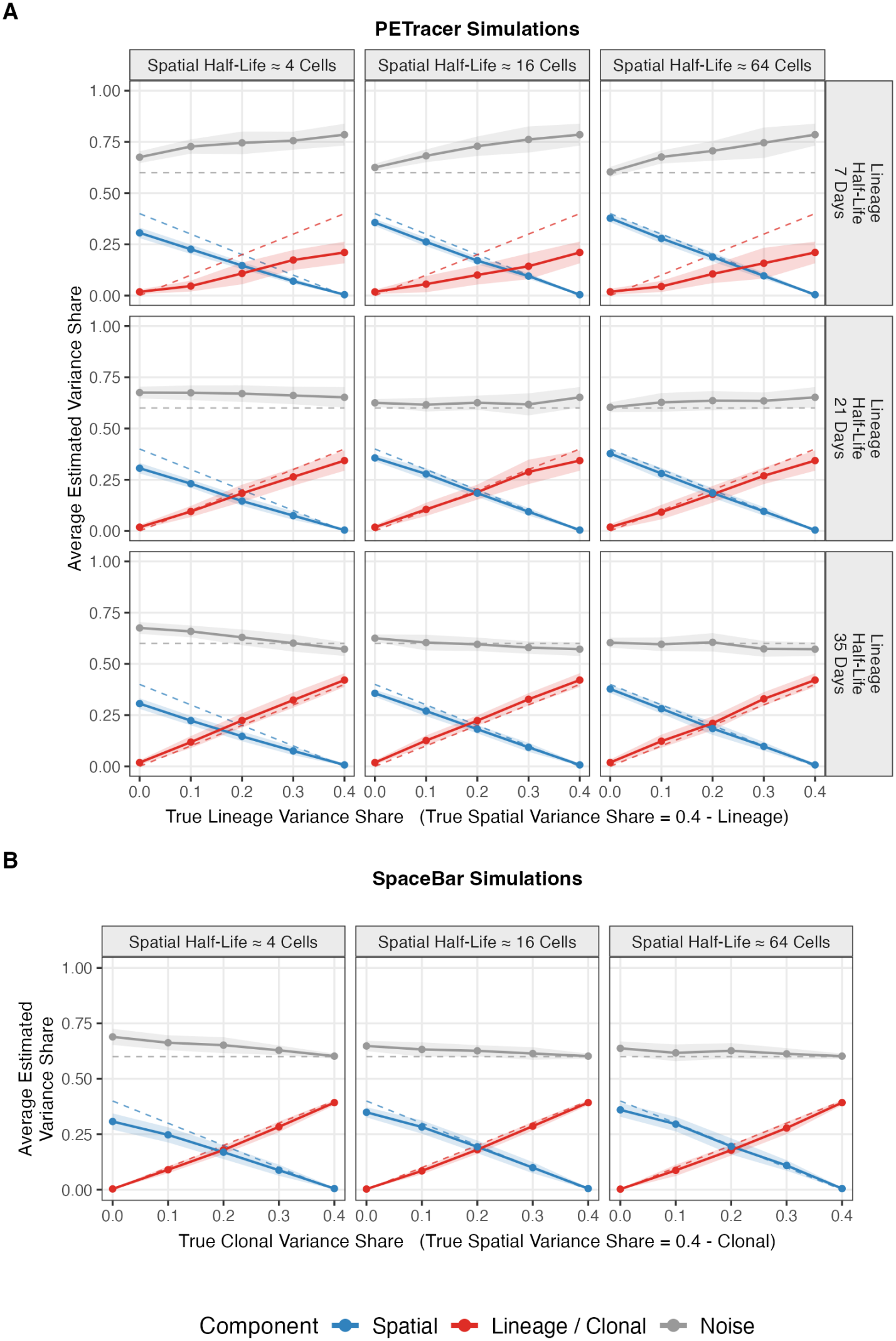
Mixed-setting PEtracer and SpaceBar simulations. Points represent average posterior mean variance shares over 25 simulations, and bands are *±* 1 standard deviation. Dashed lines indicate true variance shares. **(A)** Simulations using the lineage tree and spatial positions of PEtracer sample 3. Columns correspond to different settings of the spatial half-life, rows correspond to different settings of the lineage half-life. **(B)** Simulations using the spatial positions and clonal identities of SpaceBar sample 1.

**Fig. B7.**
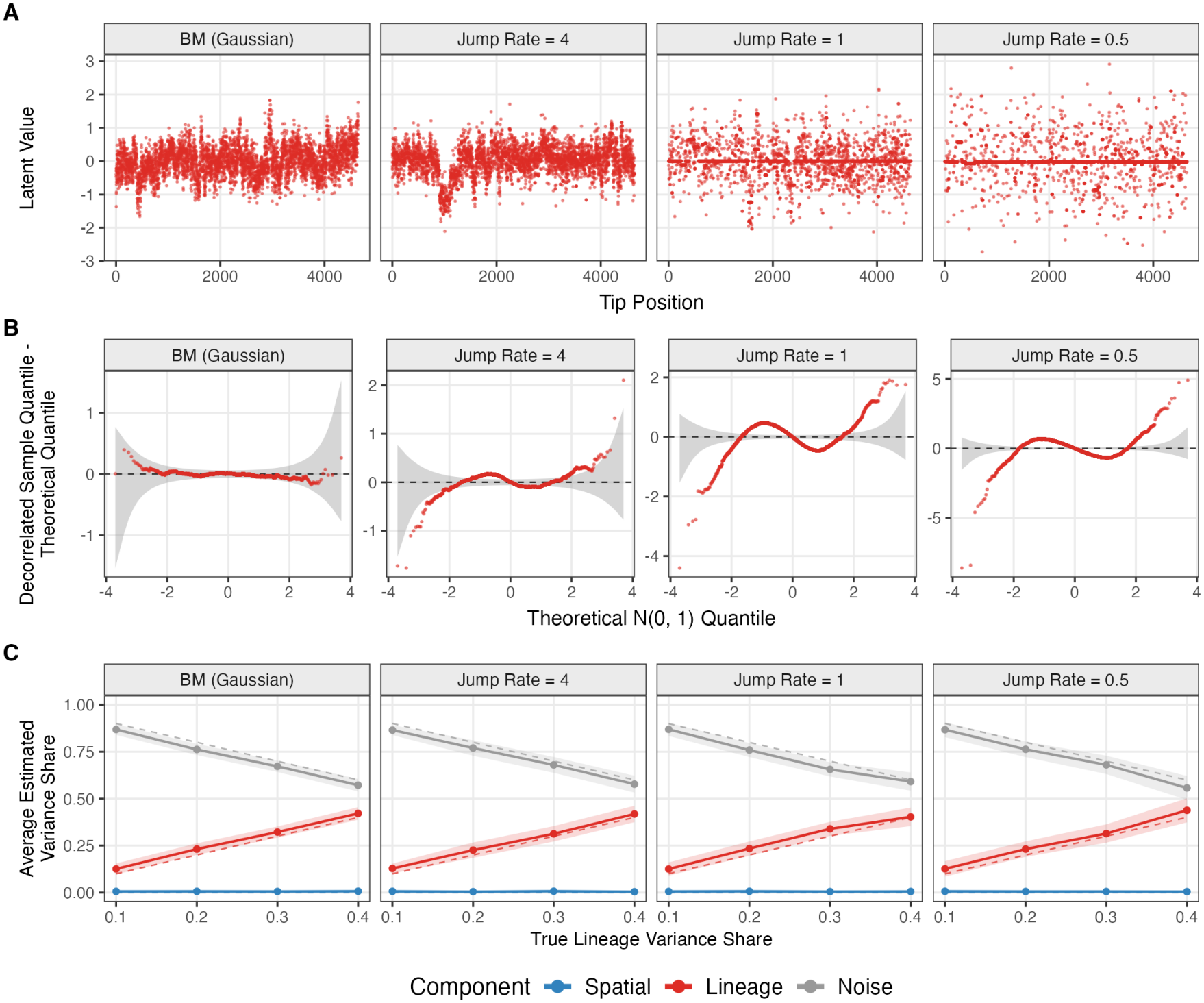
Results of Lévy misspecification simulation using the lineage tree and spatial positions of sample 3 of the PEtracer data. **(A)** An example simulated lineage field for each setting of the jump rate, defined as *ι* in Eq. B29. The x-axis is the tip position of the leaf along the lineage tree. **(B)** QQ plots of the raw lineage draws underlying (A) (same seeds, before constraining and rescaling), decorrelated using Eq. B30, with simultaneous level 0.05 bands created using qqconf [69]. **(C)** SOUR posterior mean variance shares on data generated from each arm; dashed lines are the realized true shares, bands are *±*1 SD over 25 replicates.

**Fig. B8.**
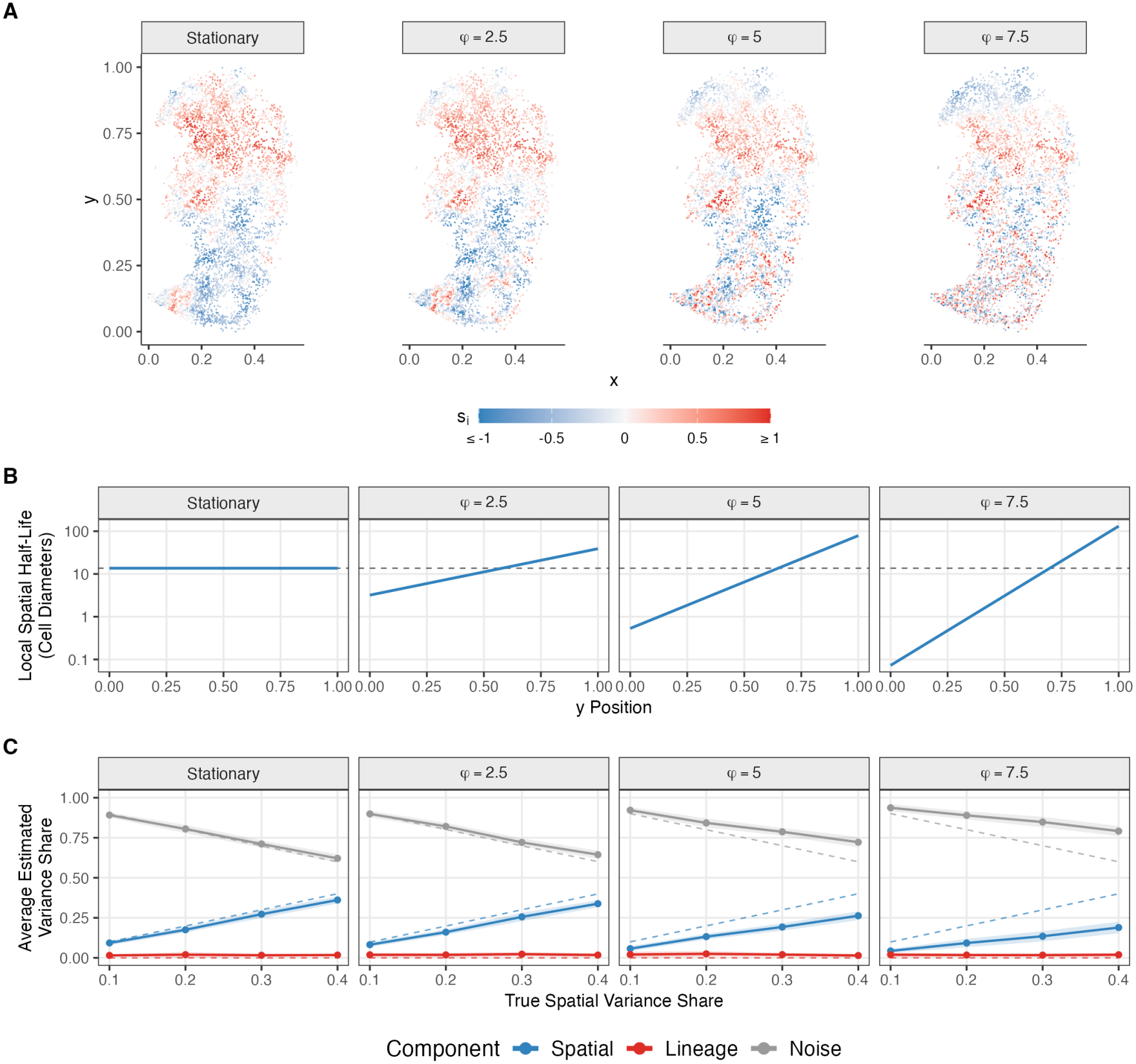
PS spatial misspecification simulation. **(A)** One example simulated spatial field per value of the gradient strength, *φ*. **(B)** The local spatial half-life against position along the gradient (y) axis. **(C)** SOUR posterior mean variance shares on data generated from each arm; dashed lines are the realized true shares, bands are *±* 1 SD over 25 replicates.

**Fig. B9.**
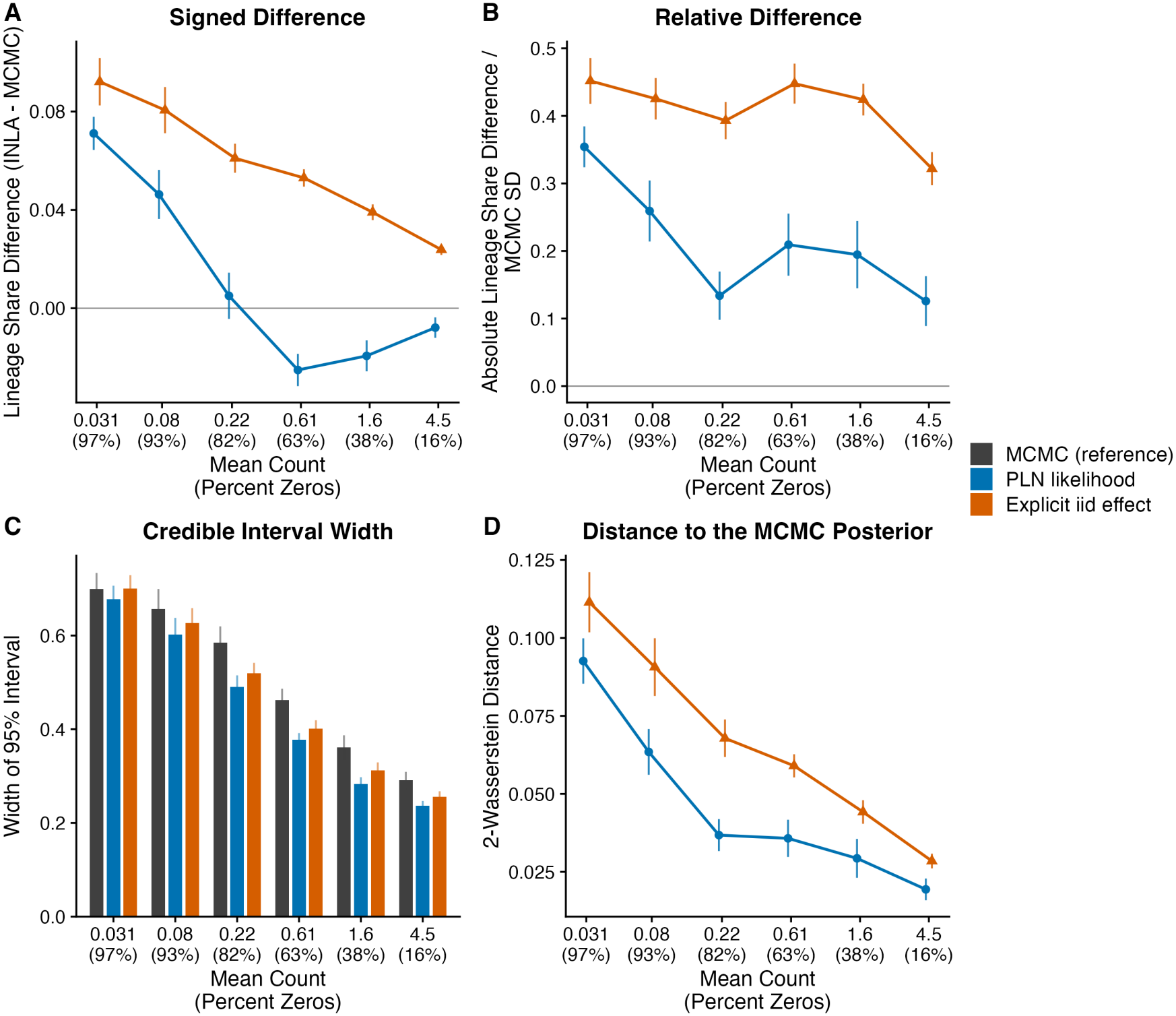
Comparison of the Poisson log-normal likelihood and an explicit iid latent effect against MCMC, on data simulated from a fixed-root OU process on a tree with 1024 leaves and a true lineage variance share of 0.5. The *x* axis of each panel gives the mean count across cells, with the corresponding percentage of zeros in parentheses. **(A)** Difference between the posterior mean lineage variance share of each INLA fit and that of the MCMC fit. **(B)** The absolute value of that difference, divided by the posterior standard deviation of the MCMC fit. **(C)** Width of the 95% credible interval for the lineage variance share, for all three fits. **(D)** 2-Wasserstein distance between the posterior of each INLA fit and the MCMC posterior. Each point or bar is the mean over 30 simulated datasets, and every interval is a 95% confidence interval for that mean.

## Appendix C Supplementary Figures

**Fig. C10.**
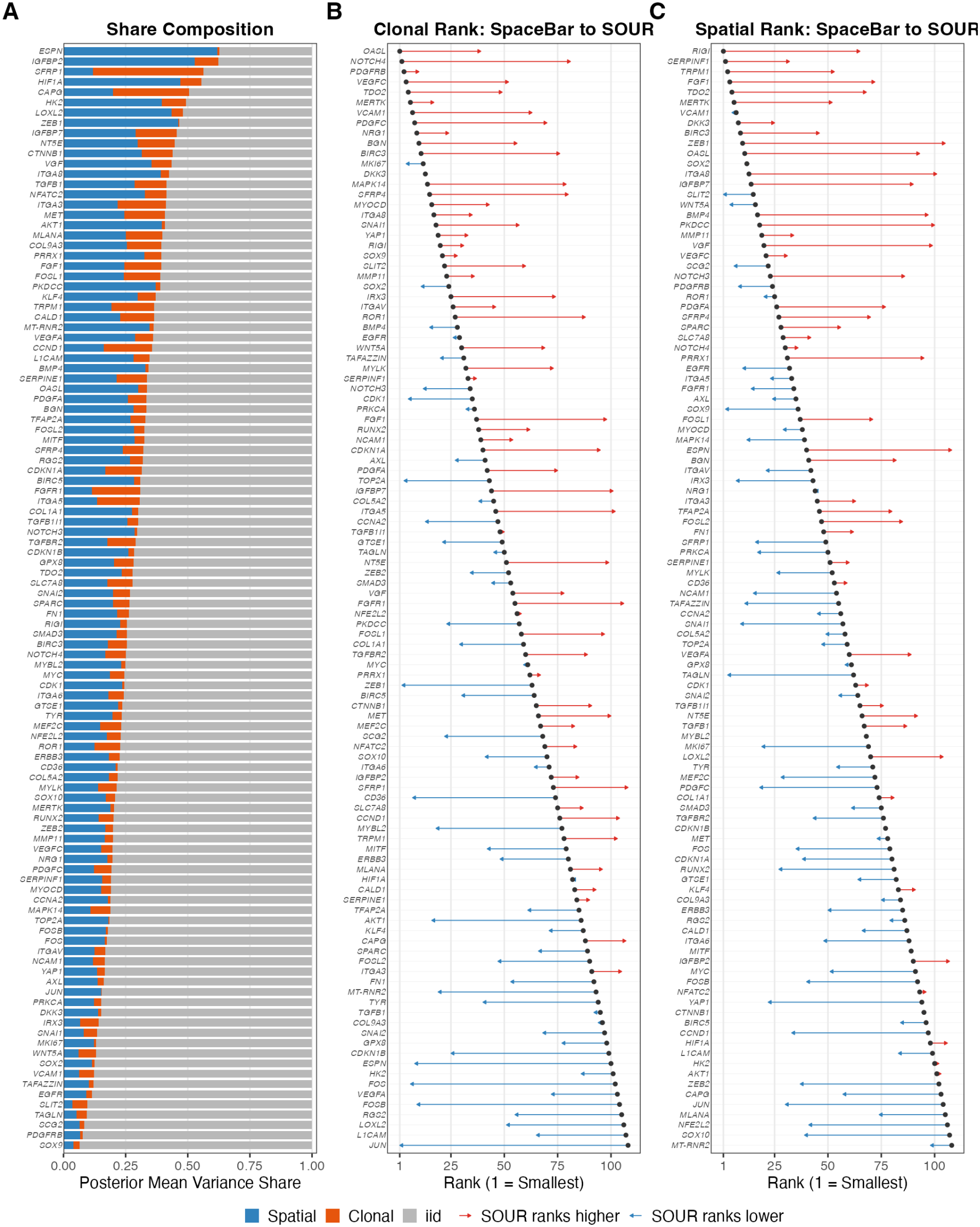
Per-gene results in sample 3 of [23]. **(A)** Posterior mean variance shares of the spatial, clonal, and noise components, ordered by total structured share. **(B)** Each gene’s rank under the marginal clone range (dot) and under SOUR’s clonal variance share (arrow head), ordered vertically by the SpaceBar rank. **(C)** As (B), for the marginal spatial rank and SOUR’s spatial share. Rank 1 is the smallest value on both axes, so a rightward arrow marks a gene SOUR ranks higher than SpaceBar.

**Fig. C11.**
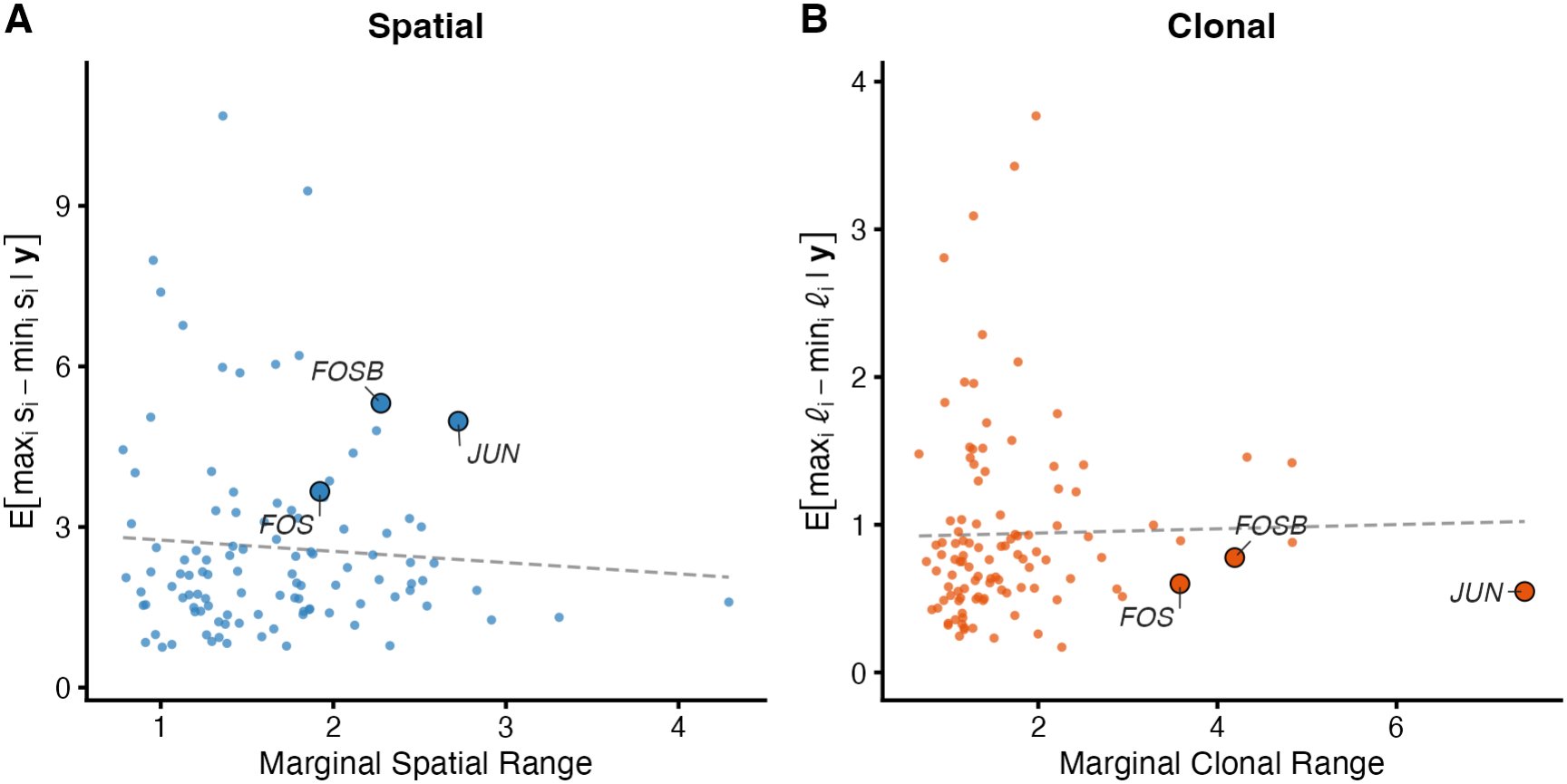
Comparison of the marginal range method, variance shares, and the posterior range of the latent fields. The horizontal axis is the published marginal range and the vertical axis is the posterior mean of the latent field range, E[max_*i*_ *u*_*i*_ *−* min_*i*_ *u*_*i*_ | **y**], for component *u*. **(A)** Spatial component. **(B)** Clonal component. Dashed lines are least squares fits.

**Fig. C12.**
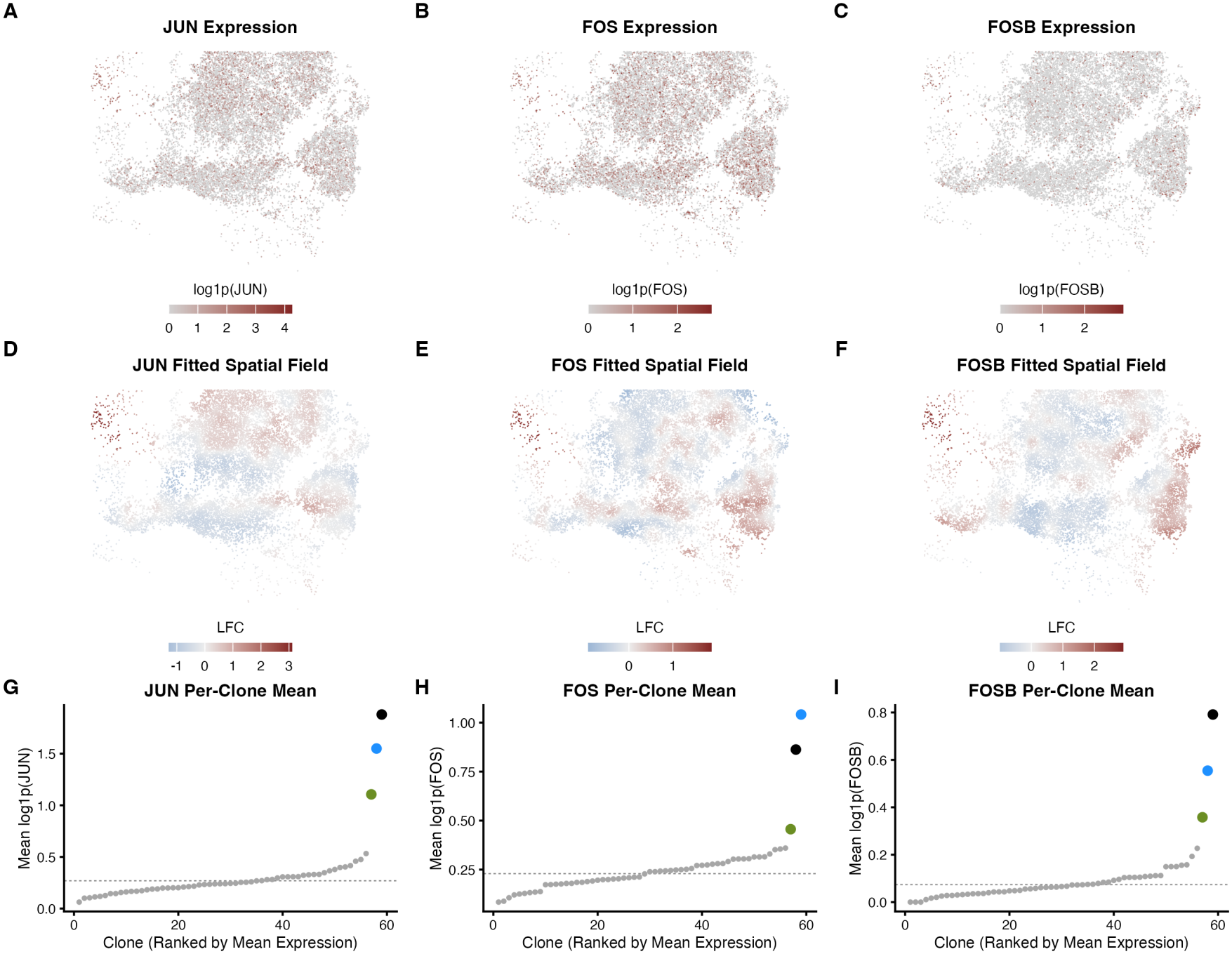
*JUN, FOS* and *FOSB* in sample 3 of [23]. **(A–C)** Expression, normalized as log(1+*y*_*i*_*/r*_*i*_). **(D–F)** Posterior mean of the spatial field **s** of the SOUR model fit to each gene. Values are a log-fold change relative to the expected expression due to the intercept *β* and lineage component ***l*. (G–I)** Mean normalized expression for each of the 59 clones, with the mean across all cells shown as a dashed line. Clone colors correspond to the top 3 clones of Fig. 4E.

**Fig. C13.**
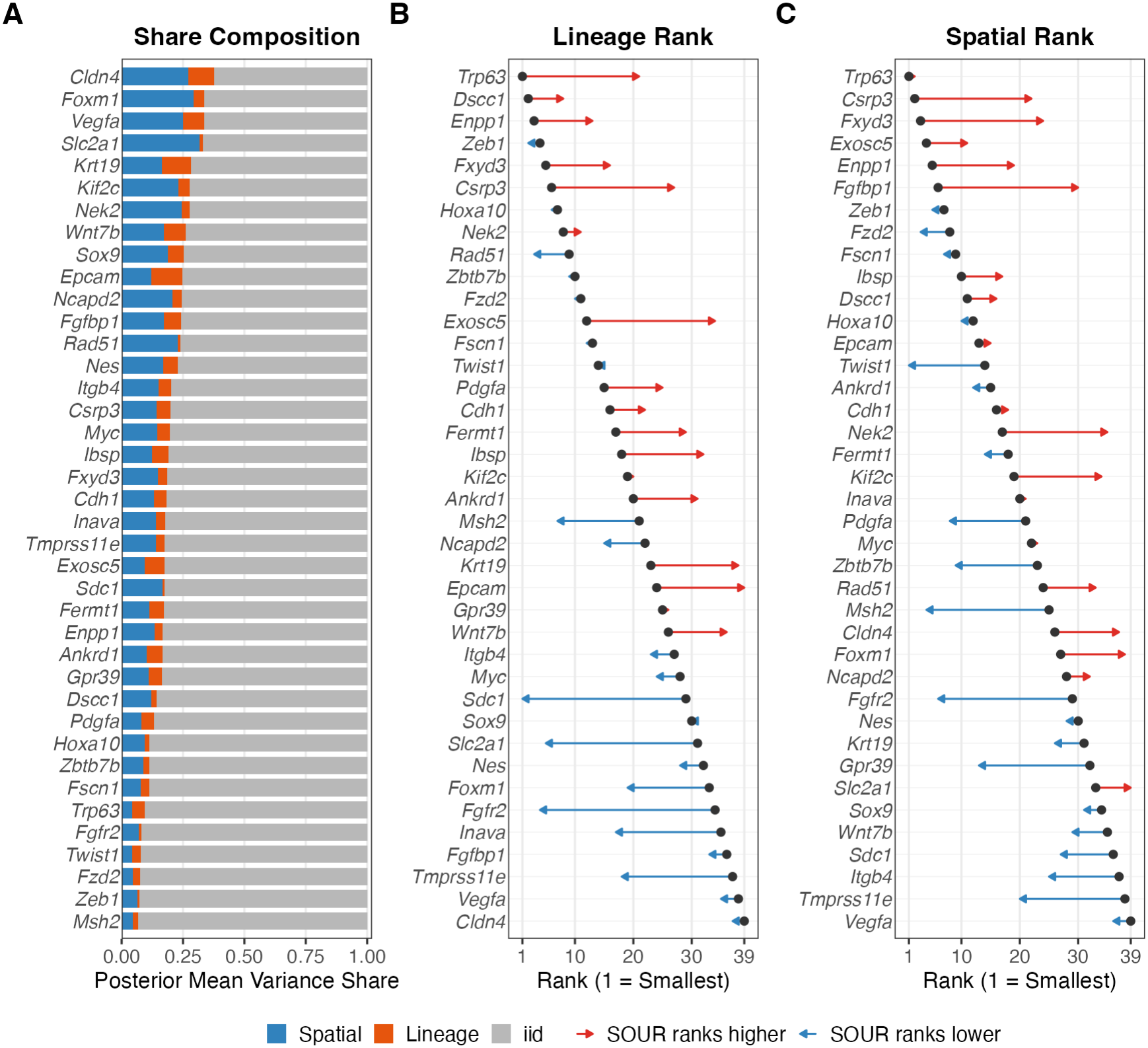
Per-gene results for the 39 genes of [21] analyzed here. **(A)** Posterior mean variance shares of the spatial, lineage, and noise components, ordered by the total share attributed to the two structured components. **(B)** Each gene’s rank under the marginal lineage autocorrelation (dot) and under SOUR’s lineage variance share (arrow head), ordered vertically by the autocorrelation rank. **(C)** As (B), for the spatial autocorrelation and SOUR’s spatial variance share. Rank 1 is the smallest value on both axes, so a rightward arrow marks a gene SOUR ranks higher than the marginal statistic does.

**Fig. C14.**
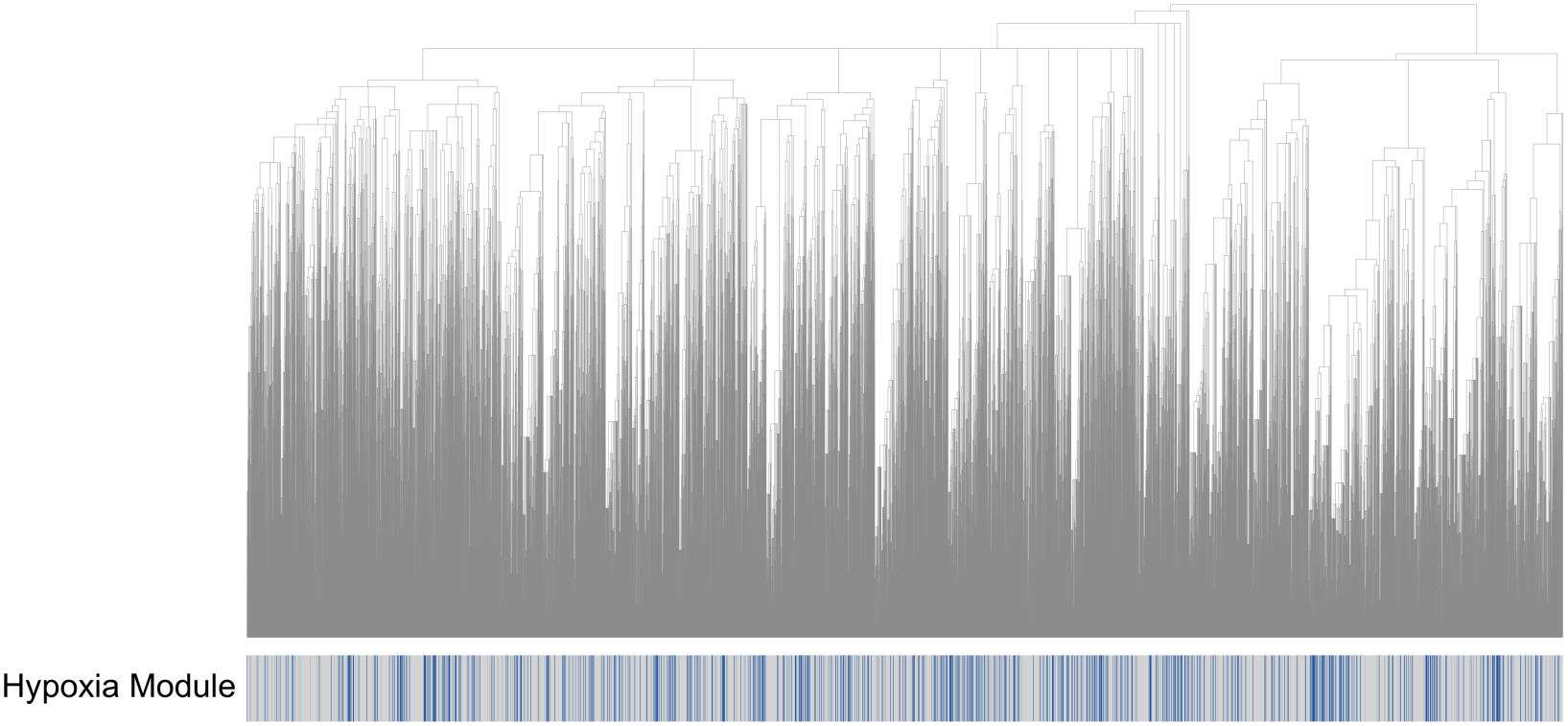
The lineage tree of the PEtracer dataset. The track beneath the tips marks, for each cell, whether it falls in the hypoxia module of [21] (blue) or not (gray).

**Fig. C15.**
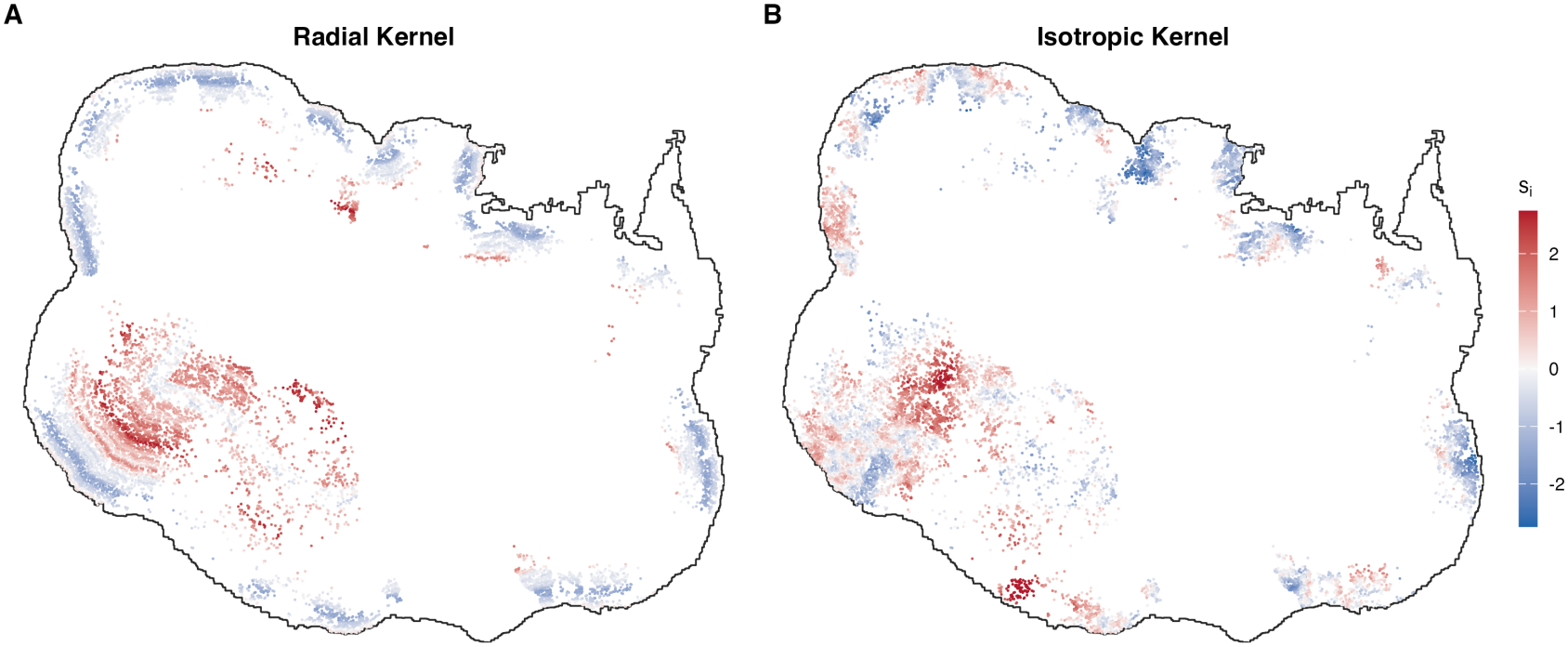
A field drawn from the radial kernel used for the SpaceBar spatial-only simulations, beside one drawn from the isotropic kernel the SOUR model assumes, on sample 1 of [23]. **(A)** One draw of 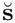 under Eq. A21, at the simulated *κ* = 0.003817 *μ*m.1, standardized to unit variance over cells. The black outline is the tumor boundary used to calculate Eq. A21. (B) One draw under the isotropic kernel of Eq. 10 at the same *κ*.

